# Radially-patterned cell behaviours during tube budding from an epithelium

**DOI:** 10.1101/308957

**Authors:** Yara E. Sanchez-Corrales, Guy B. Blanchard, Katja Röper

## Abstract

The budding of tubular organs from flat epithelial sheets is a vital morphogenetic process. Cell behaviours that drive such processes are only starting to be unraveled. Using live imaging and novel morphometric methods we show that in addition to apical constriction, radially oriented directional intercalation of placodal cells plays a major contribution to the early stages of invagination of the salivary gland tube in the *Drosophila* embryo. Extending analyses in 3D, we find that near the pit of invagination, isotropic apical constriction leads to strong cell wedging, and further from the pit cells interleave circumferentially, suggesting apically driven behaviours. Supporting this, junctional myosin is enriched in, and neighbour exchanges biased towards the circumferential orientation. In a mutant failing pit specification, neither are biased due to an inactive pit. Thus, tube budding depends on a radially polarised pattern of apical myosin leading to radially oriented 3D cell behaviours, with a close mechanical interplay between invagination and intercalation.

## Introduction

During early embryonic development, simple tissue structures are converted into complex organs through highly orchestrated morphogenetic movements. The last decade has brought a lot of understanding of individual processes and some of the important molecular players. In particular, the importance of actin together with myosin II as the major driver of cell shape changes and cell movement that underlie morphogenetic changes has been elucidated in more detail (Levayer and Lecuit, 2012; Munjal and Lecuit, 2014).

We use the formation of a simple tubular epithelial structure from a flat epithelial sheet as a model system to dissect the processes and forces that drive this change. Many important organ systems in both vertebrates and invertebrates are tubular in structure, such as lung, kidney, vasculature, digestive system and many glands. The formation of the salivary glands from an epithelial placode in the *Drosophila* embryo constitutes such a simple model of tubulogenesis (Girdler and Röper, 2014; Sidor and Röper, 2016). Each of the two placodes on the ventral side of the embryo (Fig. 1A) consists of about 100 epithelial cells, and cells in the dorso-posterior corner of the placode begin the process of tube formation through constriction of their apical surfaces, leading to the formation of an invagination pit through which all cells eventually invaginate (Fig. 1B; Girdler and Röper, 2014; Sidor and Röper, 2016).

**Figure 1.**
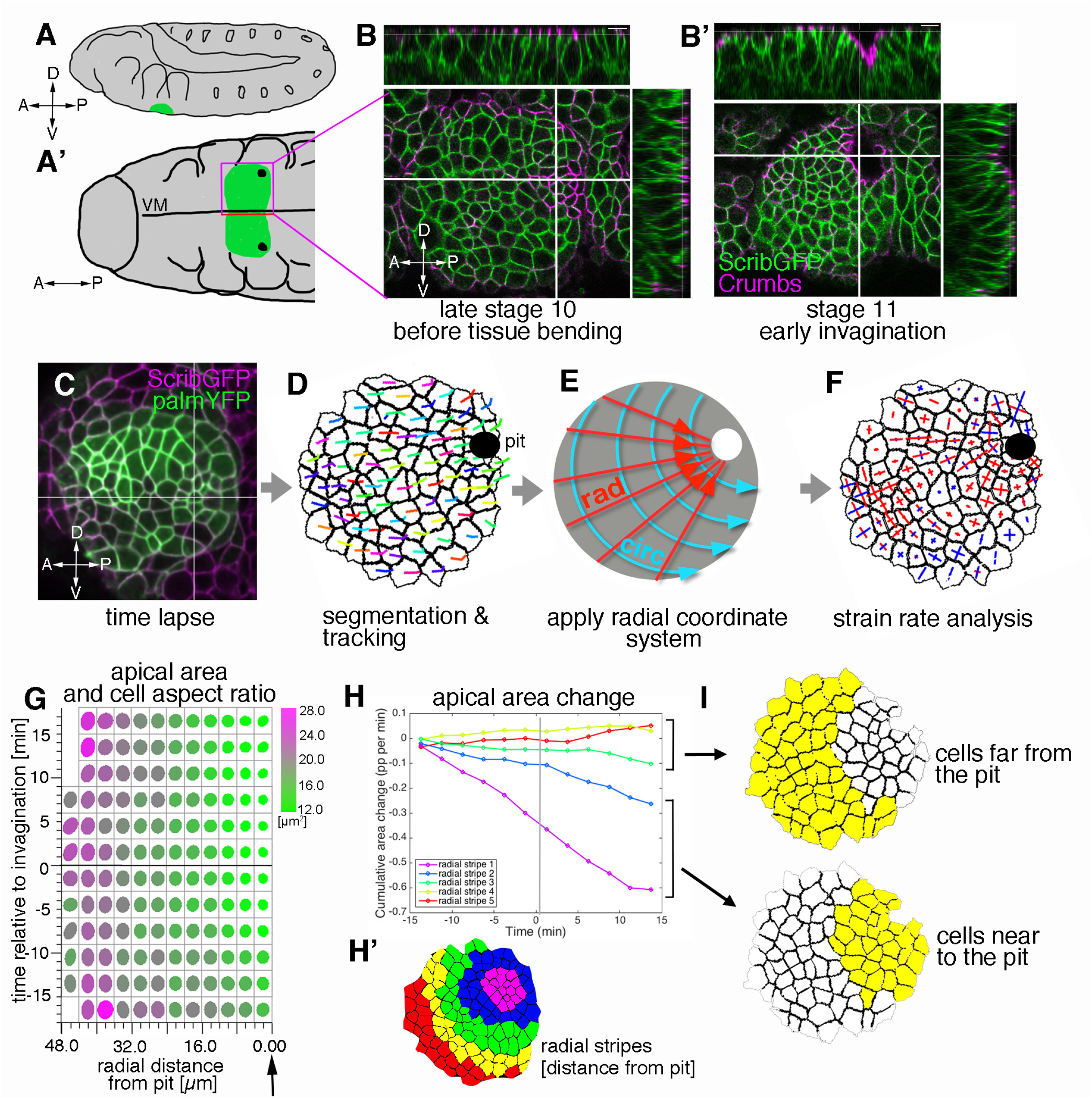
Morphogenetic events in the salivary gland placode show a radial organisation. **A, A’** Schematic of a stage 11 *Drosophila* embryo highlighting the position of the salivary gland placode (green) in lateral (**A**) and ventral (**A’**) views. **B, B’** Surface and cross-section views of the salivary gland placode just prior to the first tissue bending (**B**) and once the initial pit of invagination has formed (**B’**). Lateral membranes are labelled by ScribbleGFP (green) and apical cell outlines by Crumbs (magenta). **C-F** Workflow of the morphometric analysis. Early salivary gland placode morphogenesis is recorded by time-lapse analysis using markers of cell outlines (**C**), cells are segmented and tracked over time (**D**). Data are recorded and expressed in a radial coordinate system (**E**; ‘rad’ is the vectorial contribution radially towards the pit, ‘circ’ is the vectorial circumferential contribution). Various derived measures are projected onto the radial coordinate system. Here, projected small domain deformation (strain) rates (Blanchard et al., 2009) are shown (**F**; contraction in red, expansion in blue). See also Suppl. Movies 1 and 2. **G** Temporally and spatially resolved analysis of apical area and cell aspect ratio changes over the time interval 18 min prior to first tissue bending to 18 min after bending commences. The placode is radially organised into areas of cells with very different behaviour: cells near the invagination pit (arrow, distance 0 μm) isotropically shrink their apical area (green), whereas cells near the periphery expand anisotropically (magenta). **H-I** Cumulative time-resolved analysis of apical area change (**H**), grouping the placodal cells into radial stripes (**H’**) for the analysis. This reveals a clear split in behaviour between cells far from the pit (**I**, top;, area change value ~0) and cells near the pit (**I**, bottom; constriction, consistently negative area change)

Apical constriction, a cell behaviour of epithelial cells that can transform columnar or cuboidal cells into wedge-shaped cells and can thereby induce and assist tissue bending, has emerged as a key morphogenetic module utilised in many different events ranging from mesoderm invagination in flies, Xenopus and zebrafish to lens formation in the mouse eye (Lee and Harland, 2007; Martin and Goldstein, 2014; Martin et al., 2009; Plageman et al., 2011). Apical constriction relies on the apical accumulation of actomyosin, that when tied to junctional complexes can exert pulling forces on the cell cortex and thereby reduce apical cell radius (Blanchard et al., 2010; Mason et al., 2013). Two pools of apical actomyosin have been identified: junctional actomyosin, closely associated with apical adherens junctions, as well as apical-medial actomyosin, a highly dynamic pool underlying the free apical domain (Levayer and Lecuit, 2012; Röper, 2015).

Another prominent cell behaviour during morphogenesis in all animals is cell intercalation, the directed exchange of neighbours, that is for instance the driving force behind events such as convergence and extension of tissues during gastrulation. Also during cell intercalation apical actomyosin activity is crucial to processes such as junction shrinkage and junction extension that underlie this cell behaviour (Collinet et al., 2015; Rauzi et al., 2010). Importantly, all cell behaviours during morphogenesis require close coordination between neighbouring cells. This is achieved on the one hand through tight coupling of cells at adherens junctions, but also through coordination of actomyosin behaviour within groups of cells, often leading to seemingly supracellular actomyosin structures in the form of interlinked meshworks and cables (Blankenship et al., 2006; Röper, 2012; Röper, 2013).

The coordination between cells at the level of adherens junctions as well as actomyosin organisation and dynamics allows a further important aspect of morphogenesis to be implemented: the force propagation across cells and tissues. There is mounting evidence from different processes in *Drosophila* that force generated in one tissue can have profound effects on morphogenetic behaviour and cytoskeletal organisation in another tissue. For instance during germband extension in the fly embryo, the pulling force exerted by the invagination of the posterior midgut leads to both anisotropic cell shape changes in the germband cells (Lye et al., 2015) and also assists the junction extension during neighbour exchanges (Collinet et al., 2015). During mesoderm invagination in the fly embryo, anisotropic tension due to the elongated geometry of the embryo leads to a clear anisotropic polarisation and activity of apical actomyosin within the mesodermal cells (Chanet et al., 2017).

We have previously shown that in the salivary gland placode during early tube formation when the cells just start to invaginate, the placodal cells contain prominent junctional and apical-medial actomyosin networks (Booth et al., 2014). The highly dynamic and pulsatile apical-medial pool is important for apical constriction of the placodal cells, and constriction starts in the position of the future pit and radiates outwards from there (Booth et al., 2014). The GPCR-ligand Fog is important for apical constriction and myosin activation in different contexts in the fly (Kerridge et al., 2016; Manning et al., 2013), and *fog* expression is also clearly upregulated in the salivary gland placode downstream of two transcription factors, Fkh and Hkb (Chung et al., 2017; Myat and Andrew, 2000b). Fkh is a key factor expressed in the placode directly downstream of the homeotic factor Sex combs reduced (Scr) that itself is necessary and sufficient to induce gland fate (Myat and Andrew, 2000a; Panzer et al., 1992). *fkh* mutants fail to invaginate cells from the placode, with only a central depression within the placode forming over time (Myat and Andrew, 2000a).

Here, we use morphometric methods, in particular strain rate analysis (Blanchard et al., 2009), to quantify the changes occurring during early tube formation in the salivary gland placode. Many morphogenetic processes are aligned with the major embryonic axes of anterio-posterior and dorso-ventral. An excellent example is germband extension in the fly embryo, where polarised placement of force-generating actomyosin networks is downstream of the early anterio-posterior patterning cascade (Blankenship et al., 2006; Simoes Sde et al., 2010). In the case of the salivary gland placode, the primordium of the secretory cells that invaginate first is roughly circular, with an off-center focus due to the invagination point being located in the dorsal-posterior corner, prompting us to assess changes within a radial coordinate framework. In addition to previously characterised apical constriction (Booth et al., 2014; Chung et al., 2017), we demonstrate that radially oriented directional intercalation of placodal cells plays a major contribution to ordered invagination at early stages. We apply quantitative strain rate analysis at different apical-basal depths in the cells, and uncover cell behaviours in a 3D context: near the pit of invagination, where medial myosin II is strong (Booth et al., 2014), cells are isotropically constricting apically leading to cell wedging, and with distance from the pit cells progressively tilt towards the pit. Cells also interleave apically in a circumferential direction, i.e. they contact different neighbours along their length, a process that can be compared to a T1 transition in depth. This strongly suggests apically driven behaviours, and we show that across the placode junctional myosin II is enriched in circumferential junctions leading to polarised initiation of cell intercalation. This is followed by polarised resolution of exchanges towards the pit, thereby contributing to tissue invagination. *forkhead* mutants, that fail to form an invagination, only show unproductive intercalations that fail to resolve directionally, likely due to the lack of an active pit. Thus, tube budding depends on a radial pattern of 3D cell behaviours, that are themselves patterned by the radially polarised activity of apical myosin II pools. The continued initiation of cell intercalation but lack of polarised resolution in the *fkh* mutant, where the invagination is lost, suggest that a tissue-intrinsic mechanical interplay also contributes to successful tube budding.

## Results

### Apical cell constriction is organised in a radial pattern in the salivary gland placode

Upon specification of the placode of cells that will form the embryonic salivary gland at the end of embryonic stage 10, the first apparent change within the apical domain of placodal cells is apical constriction at the point that will form the first point of invagination or pit (Fig. 1 A,B; (Booth et al., 2014; Myat and Andrew, 2000b)). Apical constriction at this point in fact preceded actual tissue bending (Fig. 1 B). We have previously shown that apical constriction is clustered around the pit and is important for proper tissue invagination and tube formation (Booth et al., 2014). In order to discover if any further cell behaviours in addition to apical constriction drive tissue bending and tube invagination in the early placode, we employed quantitative morphometric tools to investigate the process in comparable wild-type time-lapse movies using strain rate analysis (Blanchard et al., 2009). We imaged embryos expressing a lateral plasma membrane label in all epidermal cells as well as a marker that allows identification of placodal cells (Fig.1C and Suppl.Movie 1; for genotypes see Table 1). Time-lapse movies were segmented and cells tracked using previously developed computational tools that allow for the curvature of the tissue to be taken into account (Fig. 1 C-F)(Blanchard et al., 2009; Booth et al., 2014). The cells of the salivary gland placode that will later form the secretory part of the gland are organised into a roughly circular patch of tissue prior to invagination and maintain this shape during the process (Fig.1C). Within this circular patch, the invagination pit is located at the dorsal-posterior edge rather than within the centre of the placode. We therefore employed a radial coordinate system, with the forming pit as its origin, to analyse and express any changes (Fig.1 E). We locally projected 2D strain (deformation) rates and other oriented measures onto these radial (‘rad’) and circumferential (‘circ’) axes (Fig. 1F).

**Table 1.** Genotypes used for Figure Panels.

| Figure | Panel | Embryo genotype | Number of embryos used for quantitative analysis |  |
| --- | --- | --- | --- | --- |
| 1 | B, B' | *Scribble-GFP* | panel representative of genotype | fixed |
| 1 | C | *fkhGal4,UASpalm-YFP/Scribble-GFP* | panel representative of genotype | live |
| 1 | G,H | *fkhGal4,UASpalm-YFP/Scribble-GFP and sqh[AX3]; sqh::sqhGFP42, UbiRFP-CAAX* | 1 and 8 | live |
| 2 | C,D,E | *fkhGal4,UASpalm-YFP/Scribble-GFP and sqh[AX3]; sqh::sqhGFP42, UbiRFP* | 1 and 8 | live |
| 3 | A,B | *fkhGal4,UASpalm-YFP/Scribble-GFP* | panels representative of genotype | live |
| 3 | D,E,F | *sqh[AX3]; sqh::sqhGFP42, UbiRFP-CAAX* | 5 | live |
| 4 | C,E,G | *fkhGal4,UASpalm-YFP/Scribble-GFP and sqh[AX3]; sqh::sqhGFP42, UbiRFP-CAAX* | 1 and 4 | live |
| 5 | C',C'', C''' | *fkhGal4,UASpalm-YFP/Scribble-GFP and sqh[AX3]; sqh::sqhGFP42, UbiRFP* | 1 and 6 | live |
| 5 | D,D' | *sqh[AX3]; sqh::sqhGFP42, UbiRFP-CAAX* | 3 | live |
| 5 | E,F,G | *sqh[AX3];sqh::sqhGFP42, UbiRFP-CAAX* | panels representative of genotype | live |
| 6 | A | *sqh[AX3]; sqh::sqhGFP42* | panel representative of genotype | fixed |
| 6 | B | *sqh[AX3]; sqh::sqhGFP42* | 5 | fixed |
| 6 | C | *sqh[AX3]; sqh::sqhGFP42 UbiRFP-CAAX* | panels representative of genotype | live |
| 6 | E,F | *sqh[AX3]; sqh::sqhGFP42, UbiRFP-CAAX* | 4 | live |
| 6 | G | *sqh[AX3]; sqh::sqhGFP42* | panels representative of genotype | fixed |
| 6 | H | *sqh[AX3]; sqh::sqhGFP42* | panels representative of genotype | fixed |
| 6 | I | *sqh[AX3]; sqh::sqhGFP42* | 5 | fixed |
| 7 | A | *sqh::sqhGFP42, UbiRFP-CAAX; fkh[6]* | panels representative of genotype | fixed |
| 7 | B | *sqh::sqhGFP42, UbiRFP-CAAX; fkh[6]/TM3* | panels representative of genotype | fixed |
| 7 | C | *sqh[AX3]; sqh::sqhGFP42, UbiRFP-CAAX* | panels representative of genotype | live |
| 7 | D | *sqh::sqhGFP42, UbiRFP-CAAX; fkh[6]* | panels representative of genotype | live |
| 7 | E,F (as in Fig.2 C,D,E) | *fkhGal4,UASpalm-YFP/Scribble-GFP and sqh[AX3]; sqh::sqhGFP42, UbiRFP-CAAX* | 1 and 8 | live |
| 7 | E,F | *sqh::sqhGFP42, UbiRFP; fkh[6]* | 5 | live |
| 8 | A,B | *sqh[AX3]; sqh::sqhGFP42, UbiRFP-CAAX* | 4 | live |
| 8 | A,B | *sqh::sqhGFP42, UbiRFP-CAAX; fkh[6]* | 5 | live |
| 8 | C,D | *sqh::sqhGFP42, UbiRFP-CAAX; fkh[6]* | 3 | live |
| 8 | E | *sqh[AX3];sqh::sqhGFP42 ,UbiRFP-CAAX* | 6 | live |
| 8 | E | *sqh::sqhGFP42 ,UbiRFP-CAAX; fkh[6]* | 5 | live |
| 8 | F | *fkhGal4,UASpalm-YFP/Scribble-GFP and*<br>*sqh[AX3]; sqh::sqhGFP42 ,UbiRFP* | 1 and 6 | live |
| 8 | F | *sqh::sqhGFP42 ,UbiRFP-CAAX; fkh[6]* | 5 | live |

We focused our analysis on the early stages of tissue bending and invagination, defining as t=0 the time point before the first curvature change at the point of invagination at the tissue level could be observed. Dynamic analysis of the changes in apical area and apical cell aspect ratio (a measure of cell elongation) of placodal cells, in the time interval of 18 min prior to and 18 min after the first tissue bending, revealed distinct zones of apical cell behaviour (Fig.1). Even prior to onset of tissue bending (Fig.1G, t=0), the apical area of cells near the pit (Fig.1G, black arrow and 1H) began to reduce, with cells farthest from the pit being larger and more elongated (Fig.1G). Grouping the placodal cells into ~2-cell-wide stripes concentric to the pit for apical area change analysis (Fig.1 H’) revealed a split into cells whose apical area over this time interval did not change or changed only slightly (Fig. 1H, red, yellow and green lines) and cells whose apical area progressively decreased (Fig. 1H, blue and purple lines). The clear split into two zones of differing cell behaviours, defined by radial distance from the invagination point at zero minutes (when the pit starts to invaginate), prompted us to analyse these regions independently in our strain rate analyses (Fig. 1I).

### Both apical constriction and cell intercalation contribute to the early stages of tube morphogenesis

Methods have been developed previously to calculate strain (deformation) rates for small patches of tissue, and to further decompose these into the additive contributions of the rates of cell shape change and of the continuous process of cell rearrangement (intercalation) (Blanchard, 2017; Blanchard et al., 2009) (Fig. 2A,B). The levels of both contributions vary dramatically between different morphogenetic processes (Blanchard et al., 2010; Bosveld et al., 2012; Butler et al., 2009; Etournay et al., 2015; Guirao et al., 2015). Cell divisions have ceased in the placode around the time of specification, and there is also no cell death, therefore none of these processes contributed to the overall tissue deformation in this case. Upon segmentation of cell outlines the rate of tissue deformation was calculated from the relative movement of cell centroids in small spatio-temporal domains of a central cell surrounded by its immediate neighbours and over 3 movie frames (~6 min). Independently, for the same domains, rates of cell shape change were calculated mapping best-fitted ellipses to actual cell shapes over time. The rate of cell intercalation could then be deduced as the difference between these two measures (Fig.2A,B and Suppl. Movie 2; see Material and Methods for a detailed description). The three types of strain rate measure were then projected onto our radial coordinate system.

**Figure 2.**
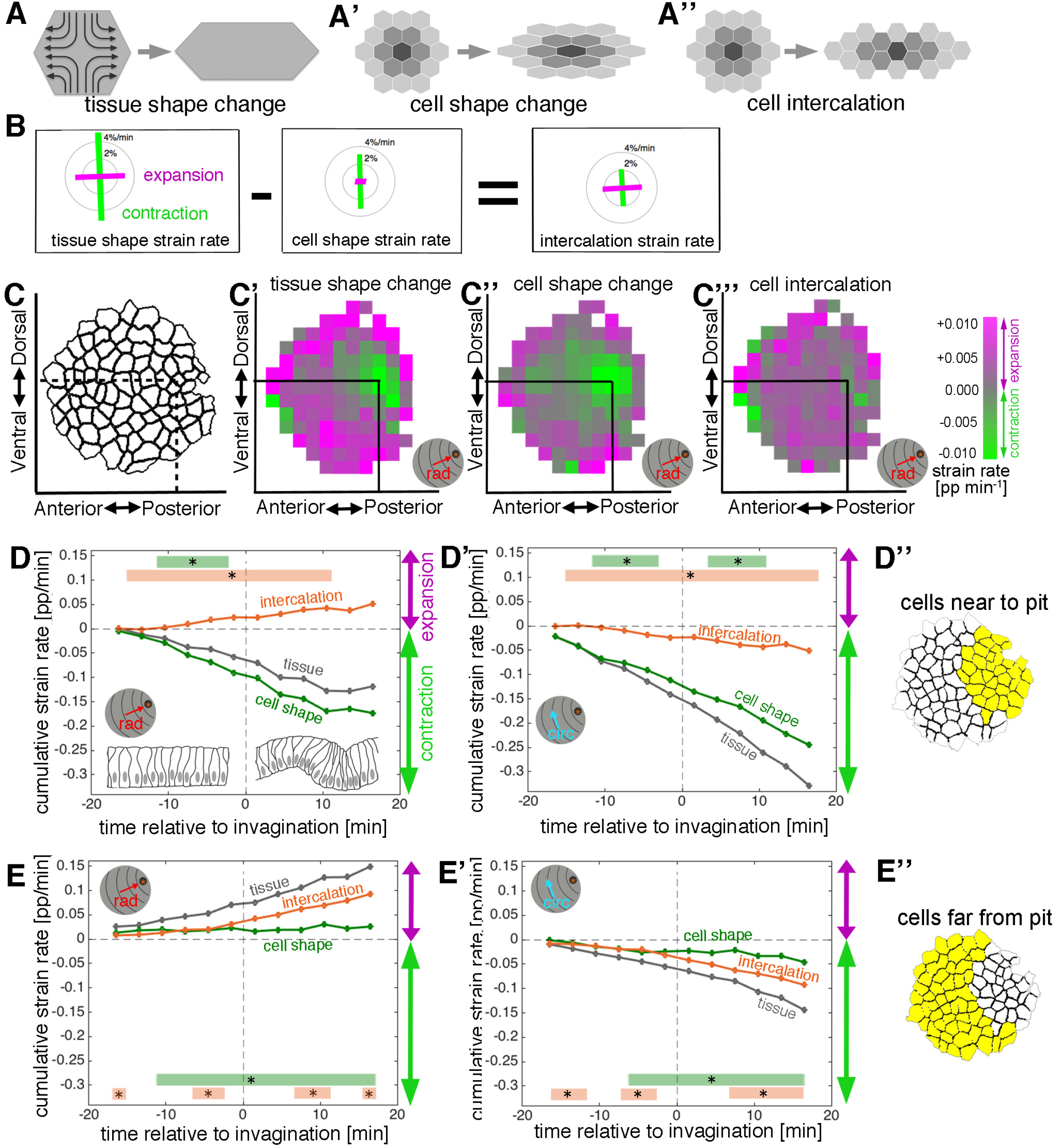
Apical strain rate analysis of early events during salivary gland placode invagination. **A-A”** During early salivary gland plcode development the change in tissue shape (**A**) can be accounted for by cell shape changes (**A’**), cell intercalation (**A”**) or any combination of the two. During this phase, cell division and gain or loss of cells from the epithelium (other than cells invaginating into the pit), do not occur. **B** For small domains of a focal cell and its immediate neighbours, tissue shape change is the sum of two additive contributions, cell shape change and cell intercalation (Blanchard et al., 2009). Both tissue shape change and cell shape change can be measured directly from the segmented and tracked time-lapse movies, so the amount of cell intercalation can be deduced. **C-C”’** Spatial depiction of the strain rate analysis covering 18 min prior to 18 min post commencement of tissue bending. Mapped onto the shape of the placode (**C**) the strain rate contribution towards the pit (‘rad’) is shown quantified from data from 9 embryos (see Suppl. Fig.2). **C’** Tissue constriction (green) dominates near the pit, with expansion (magenta) towards the periphery. The tissue constriction near the pit is mostly due to cell constriction near the pit (**C”**), whereas the tissue expansion is contributed to by both cell expansion and cell intercalation far from the pit (magenta in **C”** and **C”’**). **D-E”** Regional breakdown of time-resolved average strain rate. In the area near the pit (**D”**) tissue constriction dominates (grey curves in **D** and **D’**) and is due to isotropic constriction at the cell level (green curves in **D** and **D’**), whilst intercalation only plays a minor role in this region (orange curves in **D**, **D’**). Far from the pit (**E”**), the tissue elongates towards the pit (**E**, grey curve), with a corresponding contraction circumferentially (**E’**, grey curve), and this is predominantly due to cell intercalation (orange curves in **E** and **E’**). Statistical significance based on a mixed-effects model and a p<0.05 threshold (calculated for instantaneous strain rates [see Suppl.Fig.2]), is indicated by shaded boxes at the top of each panel: tissue vs cell shape (green) and tissue vs intercalation (orange). See also Suppl. Movie 2.

**Suppl. Figure 2.**
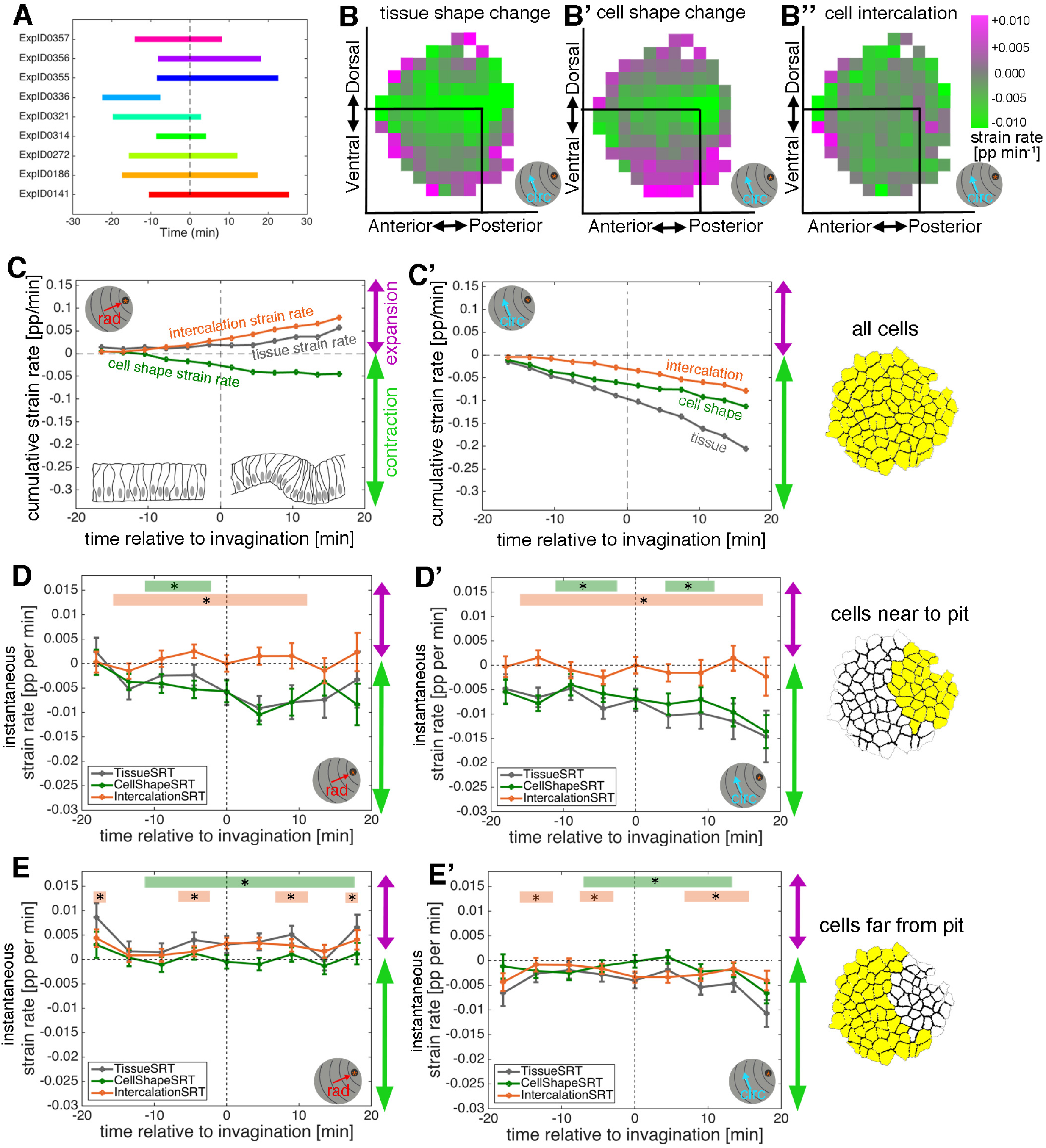
Apical strain rate analysis of early events during salivary gland placode invagination. **A** Schematic of time window covered by the time-lapse movies analysed for the apical strain rate calculations. **B-B”** Spatial summary of strain rates projected onto the circumferential orientation, covering 18 min prior to 18 min post commencement of tissue bending. **C-C’** Average strain rates of all pooled placodal cells. Overall, the tissue elongates a little radially towards the pit whilst contracting circumferentially (grey curves). Cell shapes constrict nearly isotropically (green curves), with intercalation extending the tissue towards the pit and contracting it circumferentially (orange curves). **D-E’** Instantaneous strain rates corresponding to the cumulative plots in Fig. 2 D-E”. Error bars show an indicative confidence interval of the mean, calculated as the sum of the variance of the embryo means and the mean of within-embryo variance. Statistical significance based on a mixed-effects model and a p<0.05 threshold is indicated by shaded boxes at the top of each panel (see Methods for details of statistics): tissue vs cell shape (green) and tissue vs intercalation (orange). The same conventions for displaying confidence intervals and statistical significance are used in all subsequent instantaneous strain rate plots.

Strain rate analysis of changes within the apical domain of the epithelial cells in the early salivary gland placode clearly confirmed the existence of two distinct zones of cell behaviour. Spatial plots summarising ~36 mins of data centred around the first tissue bending event (combined from the analysis of 9 wild-type embryos; Suppl.Fig.2A) revealed strong isotropic tissue contraction near the invagination pit (Fig.2C’ and Suppl.Fig.2B, green) that was mainly contributed by apical cell constriction (Fig.2C” and Suppl.Fig.2B’). At a distance from the pit, tissue elongation dominated in the radial direction (Fig.2C’, magenta) and was contributed by both cell elongation and cell intercalation (Fig. 2C”,C”’). The split in behaviour and the strong contribution of cell intercalation to the early invagination and tube formation was even more apparent from time resolved strain rate analyses. At the whole tissue level, apical cell constriction began more than 10 min before any curvature change at the tissue level (Suppl.Fig.2 C,C’, green curve), and this change was most pronounced in the cells near the invagination pit (Fig. 2 D,D’ versus E, E’, green curve). In addition, cell intercalation also commenced about 10 min prior to tissue bending (Suppl.Fig.2C,C’, orange curve), but in this case the stronger contribution came from cells far from the pit (Fig.2 E,E’ versus D,D’, orange curve).

Whereas constriction was isotropic near the pit, with equally large magnitudes contributing both radially and circumferentially (Fig. 2 D,D’, green curves), intercalation was clearly polarised towards the invagination pit, with expansion radially in the orientation of the pit (Fig. 2D,E and Suppl.Fig.2C, ‘rad’, orange curves), and contraction circumferentially (Fig. 2 D’,E’ and Suppl.Fig.2C’, ‘circ’, orange curves).

Thus, in addition to apical constriction, directional cell intercalation provides a major second cell behaviour contributing to tissue bending and invagination of a tube. Furthermore, the amount of both cell behaviours occurring was radially patterned across the placode. However, it was not clear whether these behaviours were being driven by active apical mechanisms, so we next investigated the 3D behaviours of placode cells.

### 3D tissue analysis at two depth shows coordination of cell behaviours in depth

The apical constriction as observed in the placodal cells near the pit is indicative of a redistribution of cell volume more basally, which could result in a combination of cell wedging and/or cell elongation in depth. The 3D-cell behaviour of acquiring a wedge-like shape is of particular interest in the placode because it is capable of tissue bending. Apical constriction (coupled with corresponding basal expansion if cell volume is maintained) is known to deform previously columnar or cuboidal epithelial cells into wedge-shaped cells (Martin and Goldstein, 2014; Wen et al., 2017). This is also true in the salivary gland placode, where cross-sections in xz of fixed samples and time-lapse movies confirm the change from columnar to wedged morphology (Fig.1B.B’)(Girdler and Röper, 2014; Myat and Andrew, 2000b). Most analyses of morphogenetic processes are conducted with a focus on events within the apical domain given the prevalent apical accumulation of actomyosin and junctional components (Fig.3A. A’) (Bosveld et al., 2012; Butler et al., 2009; Martin et al., 2009; Rauzi et al., 2010; Simoes Sde et al., 2010). However, this apico-centric view neglects most of the volume of the cells. For instance in the case of the salivary gland placode cells extend up to 15 μm in depth (Fig. 3B). We thus decided to analyse cell and tissue behaviour during early stages of tube formation from the placode in a 3D context. Automated cell segmentation and tracking of 3D behaviours is still unreliable in the case of tissues with a high amount of curvature, such as in the salivary gland placode once invagination begins. To circumvent this issue, we used strain rate analysis at different depth as a proxy for a full 3D analysis (Fig. 3A’,C). After accounting for the overall curvature of the tissue, we segmented and tracked placodal cells not only within the apical region (as shown in Figs 1 and 2), but also at a more basal level, ~8 μm below the apical domain (Fig. 3A’,C), and repeated the strain rate analysis (Fig. 3D-F” and Suppl. Fig.3). We used the same radial coordinate system with the pit as the origin for both layers, so we were able to compare cell behaviours between depths. Segmentation and analysis at the most-basal level of placodal cells was not reliably possible due low signal-to-noise-ratio of all analysed membrane reporters at this depth (data not shown).

**Figure 3.**
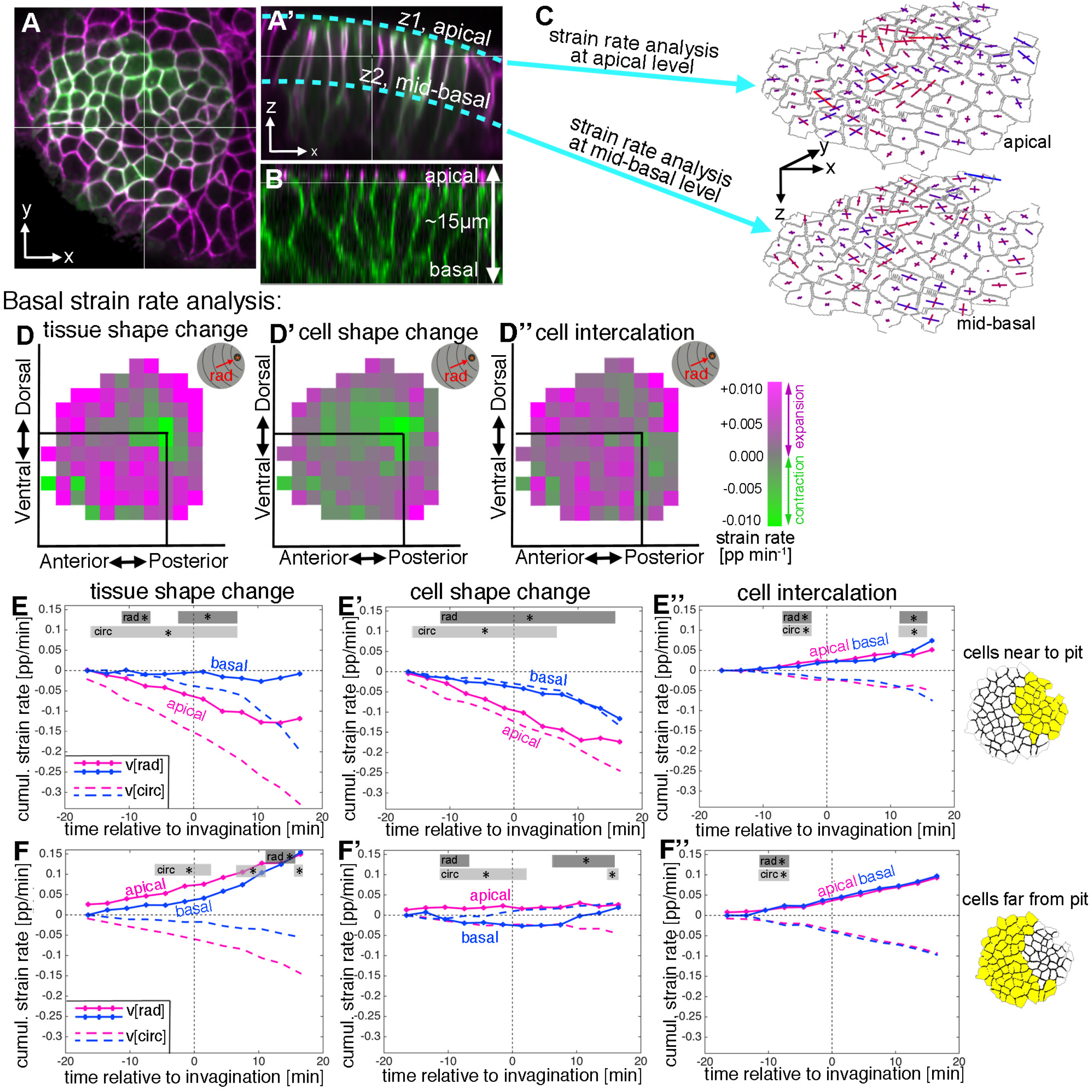
Strain rate analysis of early tubulogenesis in 3D. **A-B** Epithelial cells of the placode are about 2-5µm in apical diameter but extend about 15μm into the embryo along their apical-basal axis. To assess behaviour of the tissue and cells at depth, we analysed a mid-basal level, about 8μm basal to the apical surface. **C** We segmented and tracked cells at a depth of ~8μm, and repeated the strain rate analysis at this mid-basal level to compare with apical. **D-D”** Spatial summary of the basal strain rates (calculated from −15 to +18 min, radial contribution, “rad”). Constriction still dominates the pit region at the tissue level (**D**), due to cell constriction (**D’**), with strong expansion towards the pit everywhere else, predominantly due to cell intercalation (**D”**). **E-F”** Comparison of apical (pink, as shown in Figure 2) and basal (blue) strain rates, for tissue change (**E,F**), cell shape changes (**E’, F’**) and cell intercalation (**E”, F”**), near the pit (**E-E”**) and far from the pit (**F-F”**). Note how cell shape changes apical versus basal near the pit suggest progressive cell wedging as apices contract isotropically more rapidly than basally (**E’**), and how cell intercalation across the tissue is highly coordinated between events apically and basally (**E”, F”**). Statistical significance based on a mixed-effects model and a p<0.05 threshold (deduced from instantaneous strain rates [see Suppl.Figs 2 and 3]), is indicated by shaded boxes at the top of each panel: apical ‘rad’ vs basal ‘rad’ (dark grey) and apical ‘circ’ vs basal ‘circ’ (light grey).

**Suppl. Figure 3.**
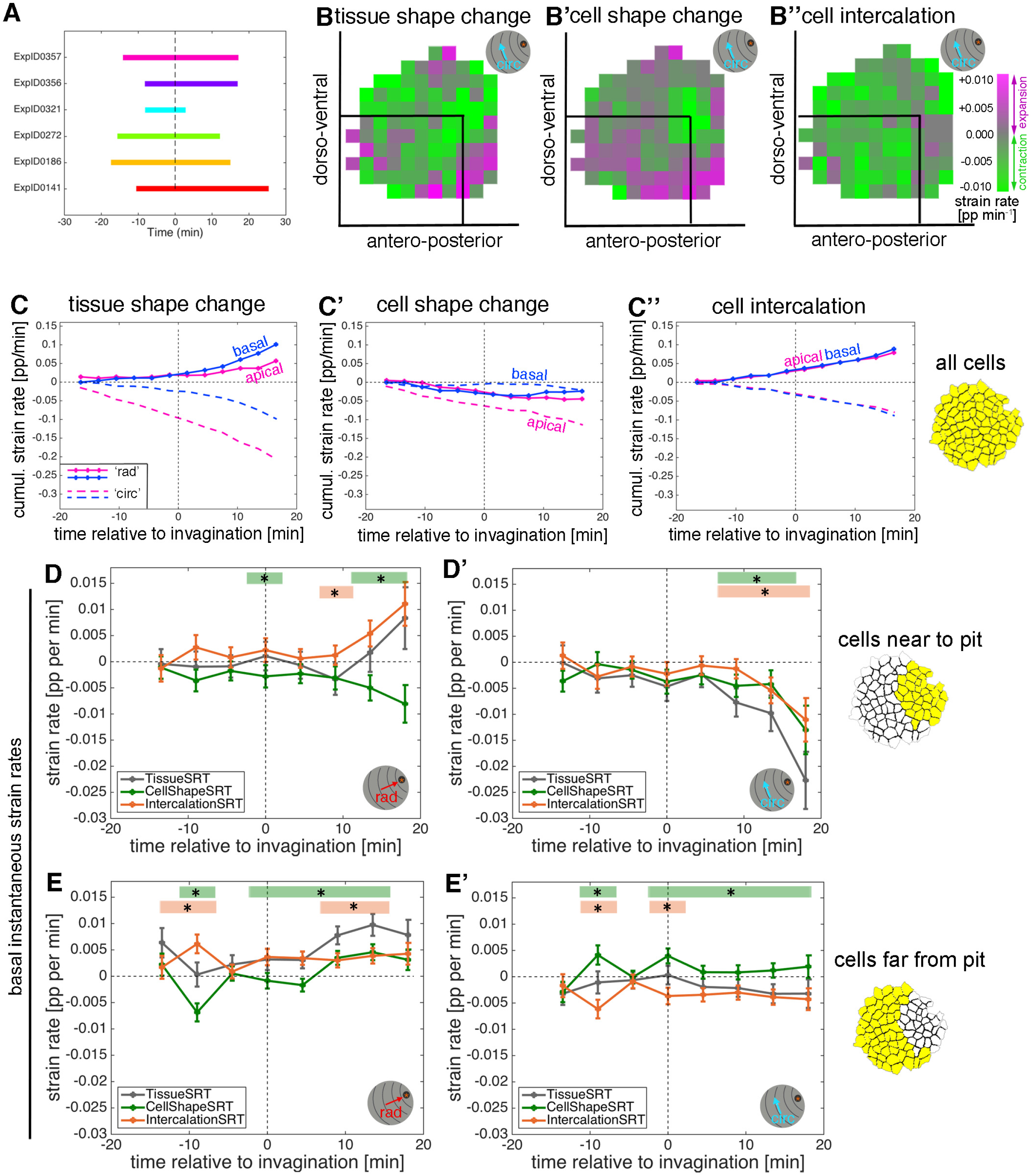
Strain rate analysis of early tubulogenesis in 3D. **A** Schematic of time window covered by the time-lapse movies analysed for the basal strain rate calculations. **B-B”** Spatial summary of basal strain rates projected onto the circumferential orientation, covering 15 min prior to 18 min post commencement of tissue bending. **C-C’** Comparison of the average basal and apical strain rates of all pooled placodal cells. Overall, basally the tissue elongates towards the pit whilst contracting circumferentially, in a more isotropic fashion than apical. Basally, cell shape change is weakly contractile radially, like apical, while it is near zero circumferentially, unlike apical (**C’**), with intercalation extending the tissue towards the pit and contracting it circumferentially almost identically in apical and basal (**C”**). **D-E’** Instantaneous basal strain rates corresponding to the cumulative plots in Fig. 3 D-E”. Error bars depict within-embryo variation of 6 movies. Significance using a mixed-effects model of p<0.05 is indicated by shaded boxes at the top of each panel: tissue vs cell shape (green) and tissue vs intercalation (orange), see Methods for details of statistics.

Overall, the spatial pattern of change at mid-basal level was similar to changes within the apical domain during 33 min centred around the start of tissue bending (Fig. 3D and Suppl. Fig.3 B). Isotropic tissue contraction at mid-basal depth was clustered around the position of the invagination pit (Fig. 3D and Suppl. Fig.3B, green), and this was due to cell constriction at this level (Fig. 3D’, and Suppl. Fig.3B’), whereas the tissue expansion at a distance from the pit in the radial direction (Fig. 3D) was due primarily to cell intercalation (Fig.3D”). Similarly to events at the apical level, this radial expansion towards the pit was accompanied by a circumferential contraction, again due primarily to cell intercalation in the region away from the pit (Suppl.Fig.3 B, B”).

Comparing apical and basal strain rates at the cell and tissue level with respect to their radial and circumferential contributions revealed an interesting picture. In temporally resolved plots, isotropic cell constriction dominated apically in cells near the pit (Fig.3E’, magenta), but with a slower rate of constriction at mid-basal depth (Fig.3E’, blue). This was confirmed by cross-section images that show that, near the pit and once tissue-bending had commenced, the basal surface of the cells was displaced even further basally than in the rest of the placode, and cells were expanded at this level, leading to an overall wedge-shape (Fig.1 B’). In cells away from the pit, similar to the apical region, cell shapes did not change much (Fig. 3 F’). In contrast to cell shape changes that diverged at depth, at least near the pit, intercalation behaviour appeared to be highly coordinated between apical and basal levels with near identical contributions at both depth across the placode (Fig. 3 E”, F” and Suppl.Fig. 3C”).

### Patterns of cell wedging, interleaving and tilt in the placode

In cells near to the pit, the faster rate of apical constriction implies that cell wedging is occurring, but we have not confirmed this in the same dataset by measuring the relative sizes of apical and basal cell diameters. Similarly, though the rates of apical and basal intercalation are remarkably similar, this does not rule out that the actual arrangement of cells at one level is ‘tipped’ ahead of the other, through interleaving in depth (akin to a T1 transition along the apical-basal axis). For example, if cell rearrangement is being driven by an active apical mechanism, we predict that apical cell contours would be intercalating ahead of basal cell contours, even while their rates remain the same. Both wedging (Fig. 4B) and interleaving (Fig.4D) have implications for the tilt, or lean, of cells relative to epithelial surface normals (Fig. 4F), with a gradient of tilt expected for constant wedging or interleaving in a flat epithelium. We therefore set out to quantify 3D wedging, interleaving and cell tilt with new methods.

**Figure 4.**
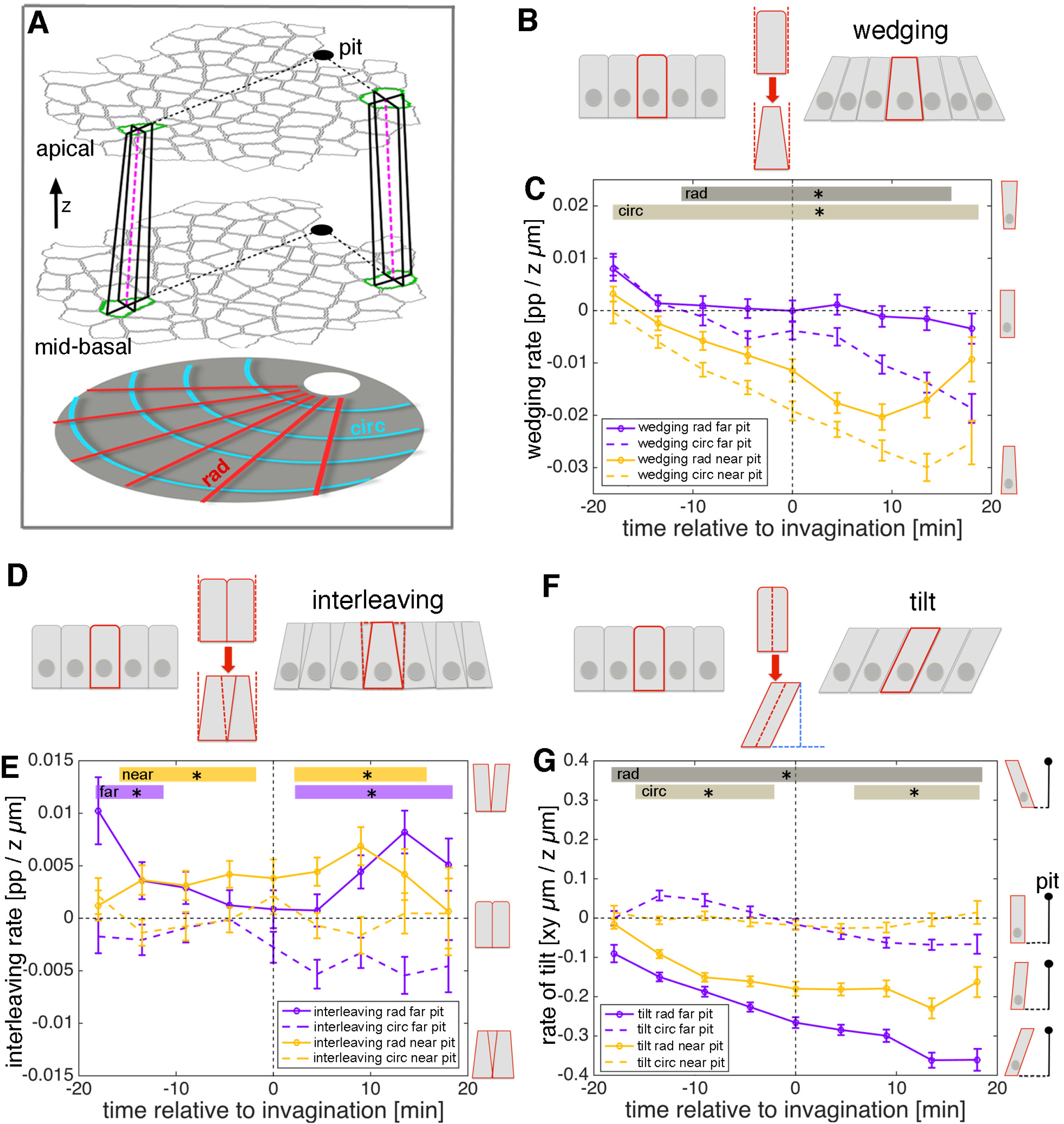
3D cell behaviours during early tube budding. **A** Calculating 3D tissue metrics using tracked cells at apical and mid-basal levels through the salivary placode. Though for simplicity only two matched cells are shown here (green outlines), all cells were matched accurately between levels (see text and Suppl. Movie 3). Small domains of a focal cell and its immediate neighbours were used to calculate local rates of wedging, interleaving and tilt (from cell ‘in-lines’, pink dashed lines) using z-strain rate methods (see text). Measures were projected onto the pit-centred radial (dotted black lines) and circumferential axes. In this tilted side-view cartoon, the z distance between apical and mid-basal levels has been exaggerated. **B** Schematic of cell wedging. **C** Cell wedging across is patterned the placode, increasing most strongly in cells near to the pit, in accordance with the isotropic apical constriction observed here (orange lines). Away from he pit, wedging contributes mainly in the circumferential direction (purple dashed line). **D** Schematic of cell interleaving. **E** Radial cell interleaving (solid line) is always more positive than circumferential interleaving (dashed lines), often significantly so. Interleaving contributes to a radial expansion apically and a concomitant circumferential contraction. **F** Schematic of cell tilt. **G** Cell tilt increased continuously in the radial direction (solid lines) apically towards the pit over the period of our analysis. A stronger rate of tilt was observed for the cells further from the pit (purple solid line), which is expected from the radial wedging seen near the pit. **C, E,G** Error bars represent within-embryo variation of 5 movies and significance at p<0.05 using a mixed-effect model is depicted as shaded boxes at the top of the panel: **C** wedging in radial orientation, near to vs far from pit (dark grey, ‘rad’) and wedging in circumferential orientation, near to vs far from pit (light grey, ‘circ). **E** Interleaving near to the pit, radial vs circumferential (orange, ‘near’) and interleaving far from the pit, radial vs circumferential (purple, ‘far’). **G** Tilt in radial orientation, near to vs far from pit (dark grey, ‘rad’) and tilt in circumferential orientation, near to vs far from pit (light grey, ‘circ).

We used the placodes for which we have tracked cells at both apical and mid-basal levels (n=5). First we developed a semi-automated method to accurately match cell identities correctly between depths (Fig. 4A and Suppl.Movie 3, and see Methods). We then borrowed ideas from recent methods developed to account for epithelial curvature in terms of the additive contributions of cell wedging and interleaving in depth (Deacon, 2012). In the early developing salivary gland placode, average tissue curvature is very slight, so we simplified the above methods for flat epithelia. We applied exactly the same methods that we have used in Figures 2 and 3 to calculate strain rates for small cell domains, but rather than quantifying rates of deformation over time, now we quantify rates of deformation in depth. The cell shape strain rate becomes a wedging rate, in units of proportional shape change per micron in z (Fig. 4B,C), the intercalation strain rate becomes the interleaving strain rate, in the same units (Fig. 4D,E), and translation velocity becomes the rate of cell tilt (Fig. 4F,G and Suppl. Fig. 4A,B; see Methods for details). Once again, we projected the z-strain rates and tilt onto our radial coordinate system.

**Suppl. Figure 4.**
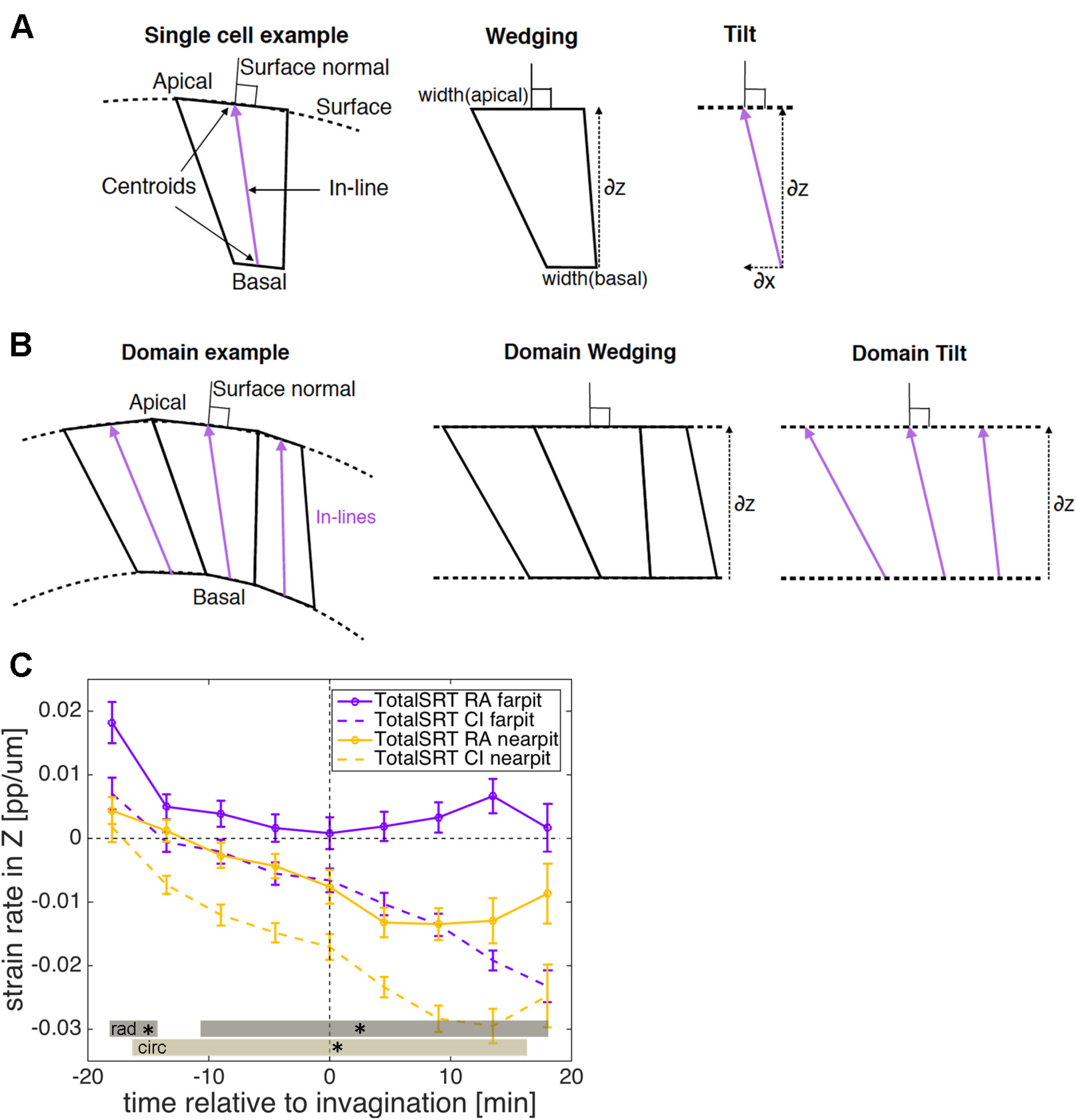
3D cell behaviours during early tube budding. **A,B** Schematics illustrating wedging and tilt definitions in the context of a single cell (**A**) and a domain of cells (**B**). **C** Total strain rate in z, representing the sum of the wedging and interleaving strain rates shown in Figure 4C, E. Error bars represent within-embryo variation of 5 movies and significance at p<0.05 using a mixed-effect model is depicted as shaded boxes at the bottom of the panel: total strain rate in z in radial orientation, near to vs far from pit (dark grey, ‘rad’) and total strain rate in z in circumferential orientation, near to vs far from pit (light grey, ‘circ).

Cells across the placode started out at −18 mins before pit invagination unwedged and mostly untilted in radial and circumferential orientations (Fig. 4C,G). Cells near the pit became progressively wedge-shaped over the next 30 minutes, with smaller apices (Fig. 4C, orange lines). Cell wedging was reasonably isotropic, but with circumferential wedging always stronger than radial. Away from the pit, progressive wedging was less rapid, again with a strong circumferential contribution but nearly no radial contribution (Fig. 4C, blue lines). That cells were less wedged radially might be because this is the orientation in which cells move into the pit, releasing radial pressure due to apical constriction near the pit.

The wedging anisotropy is also compatible with active circumferential contraction. Indeed, circumferential interleaving was always more negative than radial interleaving, often significantly so (Fig. 4E, solid vs dashed lines). Thus, interleaving contributes a circumferential tissue contraction apically, with a concomitant radial expansion. This pattern is thus also compatible with an apical circumferential contraction mechanism, possibly driving cell rearrangements.

Cell tilt, a measure of the divergence of a cell’s in-line from the surface normal (Fig. 4F and Suppl.Fig. 4A, B), increased continuously in the radial direction towards the pit over the period of our analysis (Fig. 4G, solid lines). A stronger rate of tilt was observed for the cells further from the pit (Fig. 4G’, purple solid line), which is expected from the radial wedging seen near the pit (Fig. 4C, solid orange line).

Hence, the relatively isotropic rates of apical constriction near the pit and the very similar rates of intercalation apically versus basally were in fact grounded in anisotropic wedging near the pit and in an interleaving difference between apical and basal. 3D tissue information such as wedging, interleaving and tilt are therefore essential to fully understand planar cell behaviours such as cell shape change and intercalation. Overall, our combined analysis so far suggests that isotropic apical constriction near the pit combines with an active apical circumferential contraction mechanism. We now investigate the possible origins of the latter.

### Radially polarised T1 and rosette formation and resolution underlie cell intercalation in the placode

Our strain rate analysis has revealed that the intercalation strain rate, representing the continuous process of slippage of cells past each other, was a major contributor to early tube formation and was highly coordinated between apical and basal domains. 3D domain interleaving further revealed that cell rearrangement convergence is more advanced apically in the circumferential orientation. Both these findings are measured from small groups of cells across the placode but are agnostic about neighbour exchange events or more complicated multi-neighbour exchanges. Neighbour exchanges or cell intercalations are usually thought to occur through one of two mechanisms: groups of four cells can exchange contacts through the formation of a transient 4-cell vertex structure, in a typical T1 exchange (Fig. 5A), whereas groups of more than four cells can form an intermediary structure termed a rosette, followed by resolution of the rosette to create new neighbour contacts (Fig. 5B). During convergence and extension of tissues in both vertebrates and invertebrates, the formation and resolution of these intermediate structures tend to be oriented along embryonic axis, with the resolution occurring perpendicular to the formation (Blankenship et al., 2006; Lienkamp et al., 2012). We therefore decided to also analyse intercalation in the placode by following individual events to identify the underlying mechanistic basis.

**Figure 5.**
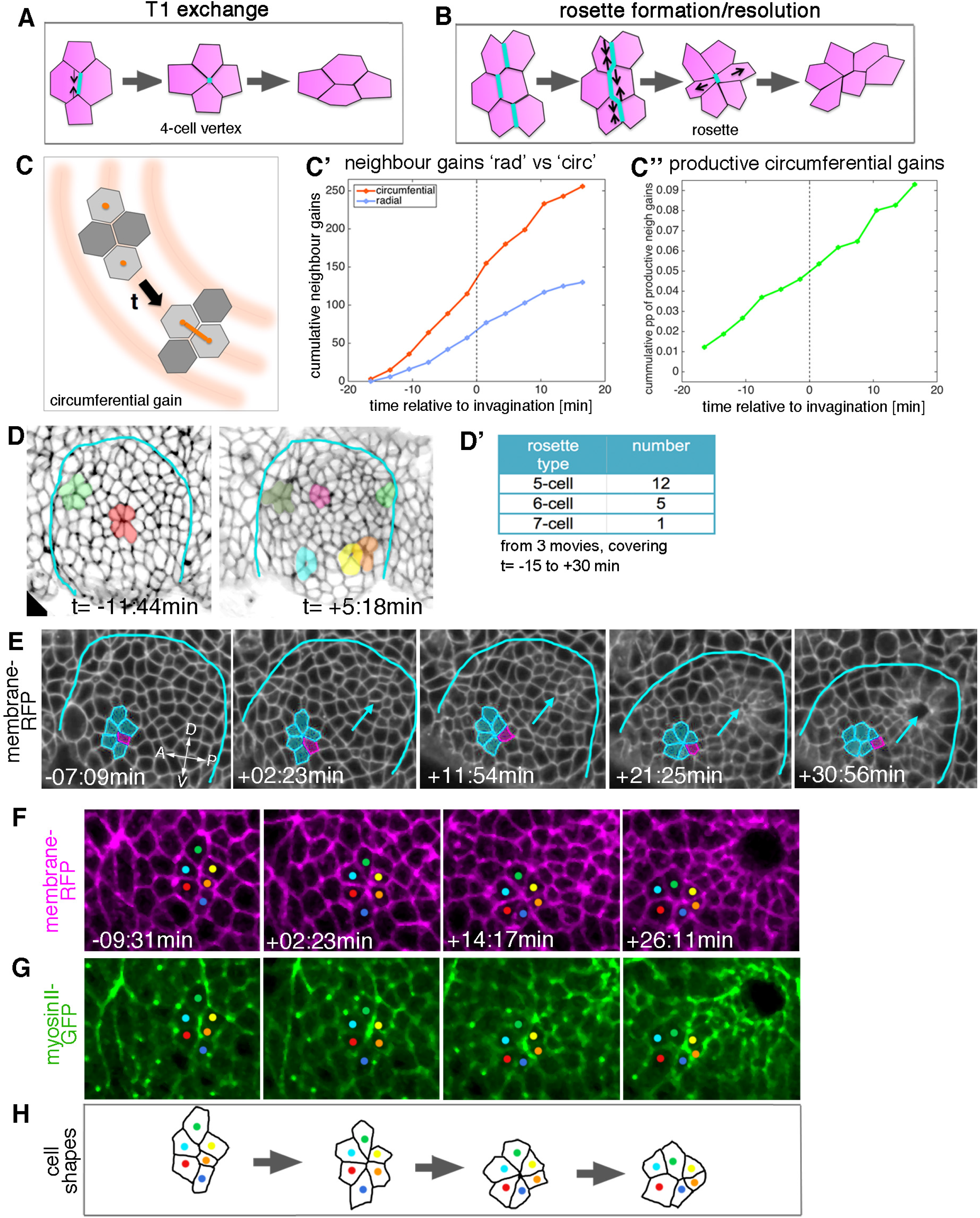
Cell intercalation during tube formation combines T1 exchanges and rosette-formation/resolution. **A,B** Depending on the number of cells involved, intercalation can proceed through formation and resolution of a 4-cell vertex, a so-called T1-exchange (**A**), or through formation and resolution of a rosette-like structure than can involve 5-10 or more cells (**B**). **C-C”** Quantification of neighbour gains as a measure of T1 and inercalation events, with an example of a circumferential neighbour gain (leading to radial tissue expansion) shown in (**C**). **C’** Circumferential neighbour gains dominate over radial neighbour gains. **C”** Rate of productive gains, defined as the amount of circumferential neighbour gains leading to radial tissue elongation and expressed as a proportion (pp) of cell-cell interfaces tracked at each time point. Data are pooled from 7 embryo movies. **D** Stills of two time-lapse movies used for the strain rate analysis, illustrating the appearance of rosette structures prior to (t= −11:44 min) and after (t= +5:18 min) tissue bending. **D’** The majority of rosettes observed in the salivary gland placode consist of 5 - 6 cells. Data are pooled from 3 embryo movies. **E-G** Stills of a time-lapse movie of an example of rosette formation-resolution. **E** The cluster of cells contracts along the circumferential direction of the placode, and resolution from the 6-cell-vertex is oriented towards the pit. Label is *Ubi-RFP-CAAX.* **F, G** The same rosette as in E in close-up, showing *Ubi-RFP-CAAX* to label membranes in magenta (**F**) and myosin II-GFP (*sqhGFP*) in green (**G**). Note the prominent but transient myosin accumulation at the centre of the rosette-forming group of cells. **H** is a schematic of the rosette formation-resolution analysed in **E-G.** See also Suppl.Movie 4.

From our database of apical cell tracks and their connectivity, we identified all T1 transitions, classifying the time point when pairs of cells become new neighbours as ‘neighbour gains’. We further sub-classified neighbour gains as being either radially or circumferentially oriented, depending on which orientation was closest to the centroid-centroid line of two cells involved in a neighbour gain (Fig. 5C). T1s occurred at a constant rate over our study period, and we observed T1s in both orientations, revealing that neighbour connectivity was quite dynamic (Fig.5C’). Nevertheless, over two-thirds of neighbour gains were oriented circumferentially, a bias that correlates with the intercalation strain rate contraction circumferentially (see for example Fig. 2E’). We defined the number of productive neighbour gains as the difference between circumferential and radial gains, since equal numbers of both would cancel each other out. In order to control for any variability between placodes or between the number of cells tracked per placode, we expressed the number of productive gains as a proportion of the number of cell-cell contacts that were available to perform a circumferential T1 per time step (see Methods). The proportion of productive circumferential gains was approximately constant, which lead to a steady net gain over time (Fig. 5C”).

In addition to typical T1 exchanges, multicellular rosette structures could easily be identified amongst the placodal cells (Fig.5 D-E). Rosette formation began prior to the first sign of tissue-bending, but the number of rosettes per placode increased afterwards (Fig. 5D,D’). Rosettes were usually formed of 5-7 cells, with most involving only 5 cells (Fig. 5D’, E). By contrast, rosettes observed during *Drosophila* germband extension can be formed of up to 12 cells (Blankenship et al., 2006). The strain rate analysis already indicated that, overall, intercalation events should be polarised to produce a contraction in the circumferential orientation with the corresponding expansion polarised towards the pit (Fig. 2 C-E’). Analysis of rosette formation and resolution in our time-lapse datasets of embryos expressing a membrane marker demonstrated that groups of cells contracted in a circumferential orientation to form a rosette, the resolution of which then moved individual cells towards the invaginating pit (Fig. 5E and Suppl.Movie 4), thereby leading to the expansion observed in the strain rate analysis.

### T1 vertex and rosette formation correlates with radial patterning of myosin and Par3/Bazooka at the apical cortex

Non-muscle myosin II is the major driver of cell shape changes in many different contexts and systems (Levayer and Lecuit, 2012; Röper, 2013; Röper, 2015), and myosin II has been shown to play an important role in T1 transitions and in rosette formation driving the convergence and extension events during germband extension in the *Drosophila* embryo (Fernandez-Gonzalez et al., 2009; Rauzi et al., 2010). We imaged embryos expressing palmitoylated RFP (*Ubi-TagRFP-CAAX*) as a membrane label and a GFP-tagged version of non-muscle myosin II regulatory light chain (called spaghetti squash, *sqh*, in *Drosophila*) under control of its own promoter in the null mutant background (*sqhAX[3]; sqhGFP42*) to assess myosin II distribution and intensity as a proxy for myosin activity (Suppl.Movie 5). In all individual rosette formation-resolution examples analysed (n= 29), junctional myosin II appeared particularly enriched in the form of short cable-like structures at the central contact sites of the rosette (Fig.5F,G). These cables initially spanned several cell diameters and shortened concomitant with the cells being drawn into a central vertex (Fig. 5H). The orientation of the short myosin cables correlated with the direction of rosette-formation, but in contrast to germband extension was not always oriented parallel to the dorsal-ventral axis, but rather following the circumferential coordinates of the placode.

When apical junctional myosin intensity was quantified in fixed samples of such *sqhGFP* embryos across the early placode, in junctions either oriented radially towards the invagination pit or circumferentially, a clear enrichment of myosin II was apparent in circumferentially compared to radially oriented junctions (Fig. 6 A, B). Such tissue-wide circumferential versus radial polarisation of myosin could assist both rosette and vertex formation during the polarised cell intercalation events.

**Figure 6.**
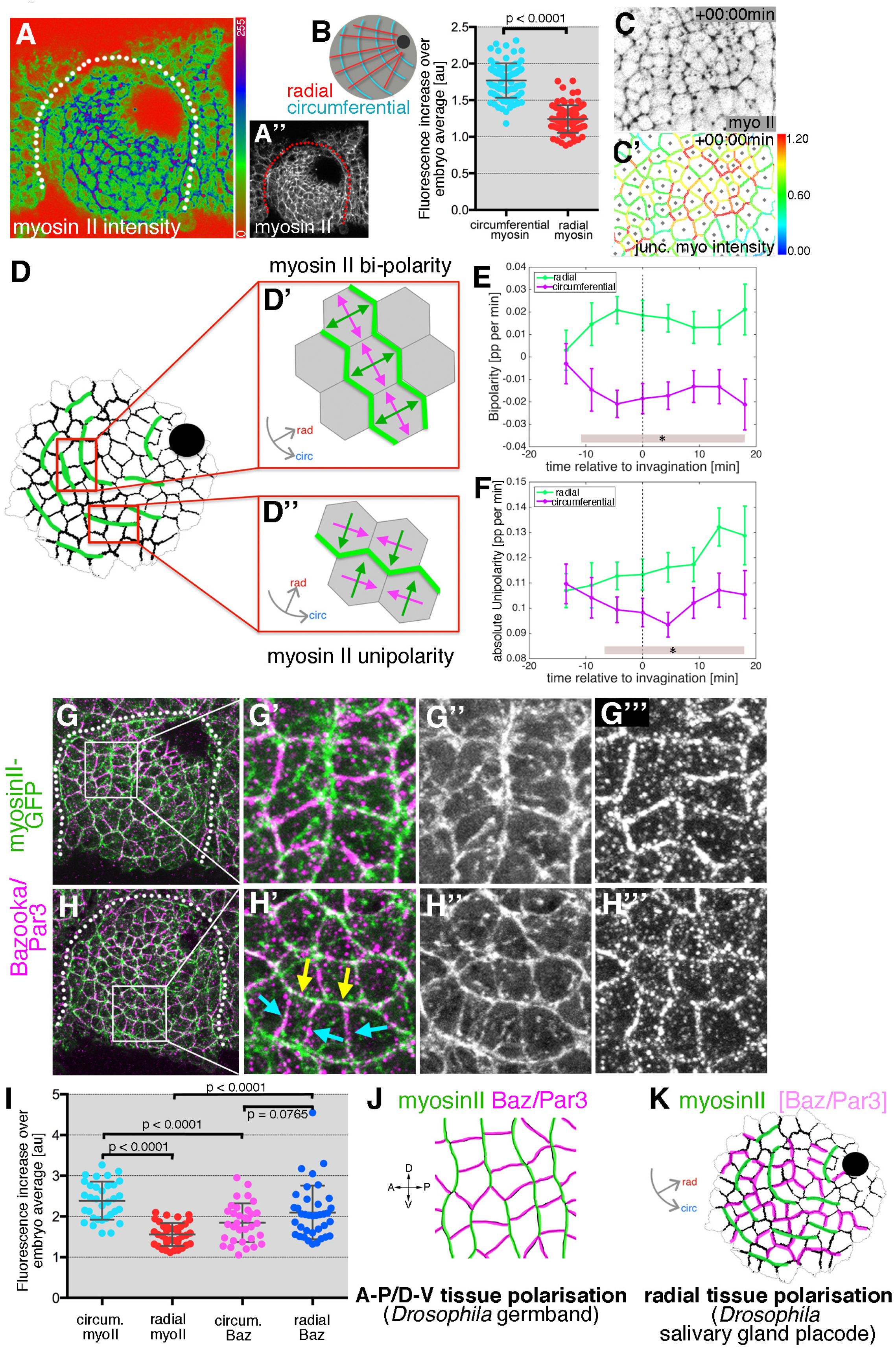
Junctional myosin II shows a strong circumferential polarisation during tube formation. **A-B** Example of junctional myosin II intensity, visualised using *sqh[AX3]; sqhGFP* in fixed embryos; **A** shows a heat map of intensity. **B** Quantification of apical junctional myosin polarisation as fluorescence increase in junctions over embryo average (5 placodes from 5 embryos, 83 circumferential junctions versus 83 radial junctions; significance calculated using unpaired t-test). **C,C’** Myosin II enrichment was quantified from segmented and tracked time-lapse movies. (**C**) depicts the sqhGFP signal from a single time point of a movie, (**C’**) shows the calculated junctional myosin II intensity. See also Suppl. Movies 5 and 6. **D-F** Myosin enrichment at junctions can occur in two flavours: (**D’,E**) Myosin II bipolarity is defined as myosin II enrichment at two parallel oriented junctions of a single cell (**D’**). Radial bi-polarity, indicating myosin II enrichment at circumferentially oriented parallel junctions (green vectors in **D’** pointing at myosin II enrichment), increases between −15 min to +18min (**E**, green curve). (**D”,F**) Myosin II unipolarity is defined as myosin II enrichment selectively on side of a cell (**D”**). Radial unipolarity, indicating enrichment at circumferentially oriented junctions (green vectors in **D”** pointing at myosin II enrichment), increases between −15 min to +18min (**F**, green curve), whereas circumferential unipolarity, indicating enrichment at radial junctions does not increase (**F**, magenta curve). **E,F** Error bars represent within-embryo variation of 4 movies, and significance between radial and circumferential bi- and unipolarity at p<0.05 using a mixed-effect model is depicted as shaded boxes at the bottom of panels E and F. See also Suppl. Movie 6. **G-H”’** Examples of myosin II and Bazooka/Par3 polarisation in fixed embryos, two different regions of two placodes are shown. Note the complementary localisation of myosin II (green) and Bazooka (magenta). **I** Quantification of myosin II and Bazooka/Par3 polarisation in areas of strong junctional myosin II enrichment at circumferential junctions as shown in **G’** and **H’**. Both circumferential (yellow arrows in **H’**) and radial (turquoise arrows in **H’**) fluorescence in crease over embryo average were quantified for myosin II and Bazooka in the same junctions (from 5 embryos; mean and SEM are shown; paired t-test for comparison in the same junctions, unpaired t-tests for comparison of circumferential myo vs radial myo and circumferential Baz vs radial Baz; N= 33 circumferential junctions and N=38 radial junctions across 5 embryos). **J,K** In contrast to the well-documented A-P/D-V polarity of cells in the *Drosophila* germband during elongation in gastrulation (**J**), the salivary gland placode appears to show a radial tissue polarity, with a radial-circumferential molecular pattern imprinted onto it that instructs the morphogenesis (**K**).

To compare and correlate junctional myosin II dynamics with the above strain rate analysis, we analysed myosin II polarisation dynamically across the whole tissue (Fig. 6C,C’). Quantitative tools that allow junctional polarisation of myosin and other players to be quantified have previously been established (Fig. 6D, D”; (Tetley et al., 2016)). Myosin II polarisation at circumferential junctions could either occur through enrichment at two opposite junctions or sides, termed bi-polarity (Fig.6D’ and Suppl.Movie 6), or through accumulation at a single junction or side within a cell, termed unipolarity (Fig. 6D” and Suppl.Movie 6). Starting about 10 min prior to tissue bending, bi-polar enrichment of myosin II at circumferential junctions was significantly stronger than at radial junctions when measured across the whole placode (Fig.6E). Similarly, starting ~5 min later, unipolar enrichment of myosin II at circumferential junctions dominated over radial enrichment when measured across the whole placode (Fig. 6F).

In cell intercalation events during germband extension, myosin polarisation is complementary to enrichment of Par3/Bazooka (Baz) as well as Armadillo/β-catenin, with both enrichments controlled by the upstream patterning and positioning of transmembrane Toll receptors and Rho-kinase (Rok) (Blankenship et al., 2006; Pare et al., 2014; Simoes Sde et al., 2010). We therefore analysed whether such complementarity was also present in the early salivary gland placode. Antibody labelling of Baz often showed a complementarity in membrane enrichment to myosin II (Fig. 6 G-I), most pronounced where myosin II was organised into circumferential mini cables during intercalation events (Fig.6G,H). Baz was enriched at radially oriented junctions where myosin II was low, and vice versa at circumferential junctions, though in contrast to germband extension, the Baz polarisation did not extend uniformly across the tissue and was overall less strong than the myosin II polarisation (Fig.6I).

Thus, in order to adapt to a circular tissue geometry and to the need for ordered invagination through a focal point during the process of tube budding, a conserved molecular pattern of myosin-Baz complementarity is apparently imprinted onto the salivary gland placode in a radial coordinate pattern, rather than the prevailing A-P/D-V pattern of the earlier embryo during germband extension (Fig.6J,K).

### Resolution but not initiation of cell intercalation is disrupted in the absence of Forkhead

In order to address how the radial pattern of behaviours and molecular factors across the placode is established, we analysed mutants in a key factor of salivary gland tube invagination, the transcription factor Fork head (Fkh). Fkh is expressed just upon specification of the placodal cells, directly downstream of the homeotic factor Scr (Zhou et al., 2001). In *fkh* mutants, invagination of the placode fails, and towards the end of embryogenesis salivary gland-fated cells undergo apoptosis as Fkh appears to prevent activation of pro-apoptotic factors (Jurgens and Weigel, 1988; Myat and Andrew, 2000a). Previous studies have concluded that Fkh promotes cell shape changes important for invagination, in particular apical constriction (Chung et al., 2017; Myat and Andrew, 2000a). Fkh is not the only transcription factor important for correct changes during invagination, but works in parallel to e.g. Huckebein (Hkb), the lack of which also confers significant problems with apical cell shape changes and invagination (Myat and Andrew, 2000b; Myat and Andrew, 2002).

We combined a *fkh* null mutant (*fkh[6]*) with markers allowing membrane labeling and cell segmentation (*Ubi-TagRFP-CAAX*) as well as myosin II quantification (*sqhGFP*; see Suppl.Movie 7)). When *fkh* mutant placodes were compared to wild-type ones at late stage 11 (beyond the time window analysed here) then *fkh* mutant placodes showed no sign of a dorsal-posterior invagination point (Fig.7 A-B’). In fact, many placodes beyond late stage 11 showed a centrally located shallow depression (Fig. 7A,A’, yellow dotted lines). Strain rate analysis of 5 segmented movies of *fkh[6]* mutant placodes, spanning the equivalent time period to the wild-type movies analysed above (Suppl. Fig.7C), showed that there was no constriction at the tissue level near the pit (Fig. 7C-E and Suppl. Fig. 7 D, D’). In fact if anything there was a slight tissue expansion (Fig.7E) caused by a slight expansion at the cell level (Fig. 7E’), with zero intercalation (Fig. 7 E”). Away from the pit *fkh[6]* mutant placodes expanded slightly (Fig.7F and Suppl. Fig.7 E, E’), again mostly due to cell shape changes (Fig. 7F’, F”).

**Figure 7.**
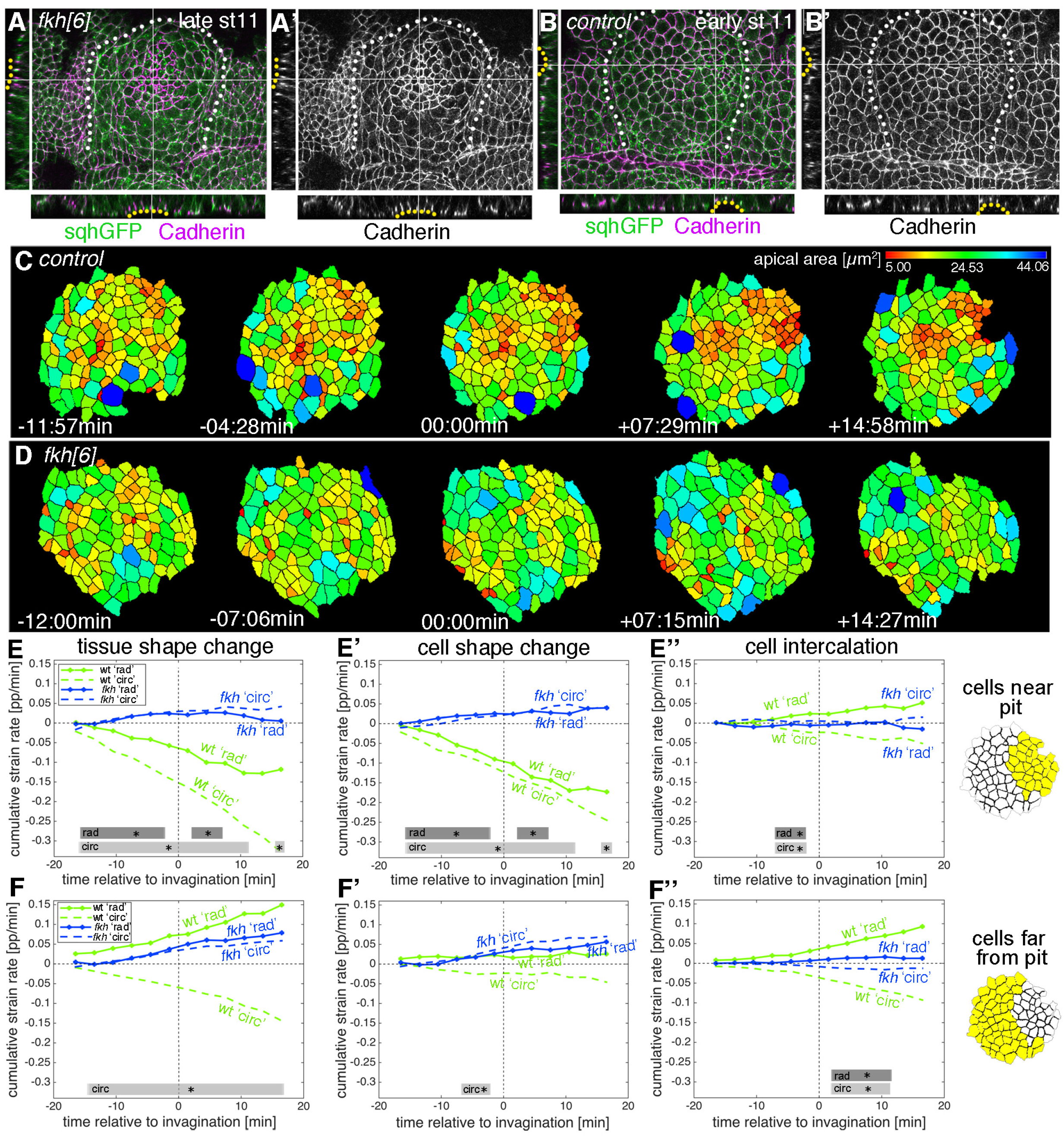
Loss of radially polarisation of cell behaviours in salivary gland placodes lacking Fkh. **A-B’** Examples of *fkh* mutant and wild-type placodes illustrating the lack of pit formation. A control placode at early stage 11 shows clear constriction and pit formation in the dorsal-posterior corner (yellow dotted line in cross-sections in **B, B’**), whereas even at late stage 11 a *fkh* mutant placode (beyond the time frame of our quantitative analysis) does not show a pit in the dorsal-posterior corner, instead a shallow central depression forms with some constricted central apices (yellow dotted line in cross-sections in **A, A’**). Myosin II (sqhGFP) is in green and DE-Cadherin in magenta, white dotted lines in main panels outline the placode boundary. **C,D** Stills of segmented and tracked time lapse movies for control (**C**) and *fkh[6]* mutant (**D**) placodes, apical cell outlines are shown and colour-coded by cell area. Note the lack of contraction at the tissue and cell level in the *fkh[6]* mutant. **E-F”** Apical strain rate analysis of *fkh* mutant placodes (from 5 movies) in comparison to the analysis of wild-type embryos (as shown in Figure 2). Over the first 36 min of tube budding centred around the first appearance of tissue-bending in the wild-type and an equivalent time point in the *fkh* mutants, the *fkh* mutant placodes show only a slight expansion at the tissue level in the cells far from the predicted pit position (**F**) due to cell shape changes (**F’**) with very little intercalation contributing to the change (**E”, F”**). Statistical significance based on a mixed-effects model and a p<0.05 threshold (calculated for instantaneous strain rates [see Suppl.Fig.2 and 7]), is indicated by shaded boxes at the top of each panel: wt ‘rad’ vs *fkh* ‘rad’ (dark grey) and wt ‘circ’ vs *fkh* ‘circ’ (light grey).

**Suppl. Figure 7.**
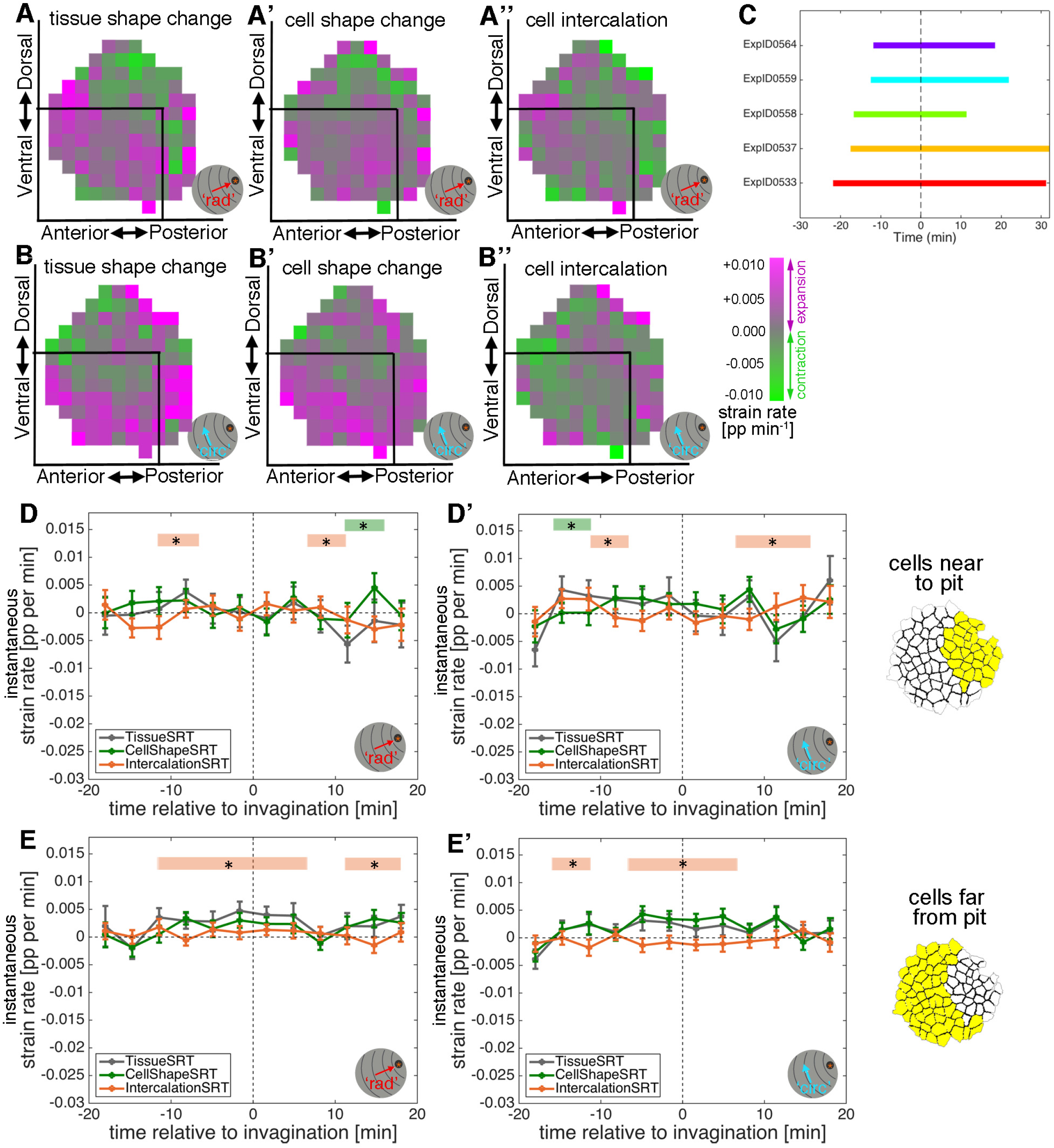
Loss of radially polarisation of cell behaviours in salivary gland placodes lacking Fkh. **A-B”** Spatial summary of average apical strain rates in *fkh[6]* mutants covering 15 min prior to 18 min post commencement of tissue bending, showing the radial (**A-A”**) as well as circumferential (**B-B”**) contributions. **C** Schematic of time window covered by the time-lapse movies analysed for the apical strain rate calculations in *fkh[6]* mutant embryos. **D-E’** Instantaneous apical strain rates in *fkh[6]* mutants corresponding to the cumulative plots in Fig. 7 E-F”. Error bars depict within-embryo variation of 5 movies. Significance using a mixed-effects model of p<0.05 is indicated by shaded boxes at the top of each panel: tissue vs cell shape (green) and tissue vs intercalation (orange), see Methods for details of statistics.

Is this strong reduction of cell behaviours due to a complete ‘freezing’ of the placode in the *fkh[6]* mutant? We analysed whether junctional myosin II was still polarised in the *fkh[6]* mutant placodes. In fixed and live samples, *fkh[6]* mutant placodes still showed the circumferential actomyosin cable surrounding the placode (Suppl.Fig.8E’ and I”’, compare to D’ and H”’; (Röper, 2012)). When myosin II unipolarity and bi-polarity were quantified from 5 segmented and tracked movies, at the tissue-level there was a strong and significant reduction in myosin II polarisation, with no difference between radial and circumferential orientations in either measure and myosin II bi-polarity indistinguishable from zero (Fig. 8 A,B). Nonetheless, cells were not static but in fact neighbour exchanges were present, both in form of T1 exchanges and rosettes (Fig.8C-F). The quantitative analysis of neighbour gains from segmented and tracked movies in the *fkh[6]* mutant revealed that gains accumulated both circumferentially as well as radially, at a rate comparable to that observed in the control (Fig. 8E). However, in contrast to the control, where circumferential gains significantly outweighed radial neighbour gains leading to a positive net rate of productive circumferential gains (Fig.8F, green line), in the *fkh[6]* mutant circumferential and radial gains occurred in equal amounts (Fig.8E) leading to a near zero net gain (Fig. 8F, pink line). Interestingly, when focusing on individual events such as rosettes, despite a loss of tissue-wide myosin II polarisation, myosin II was still enriched at the central constricting junction (Fig. 8Cd’), and in still pictures remnants of myosin II-Baz complementarity could be identified (Suppl.Fig.8L-M).

**Figure 8.**
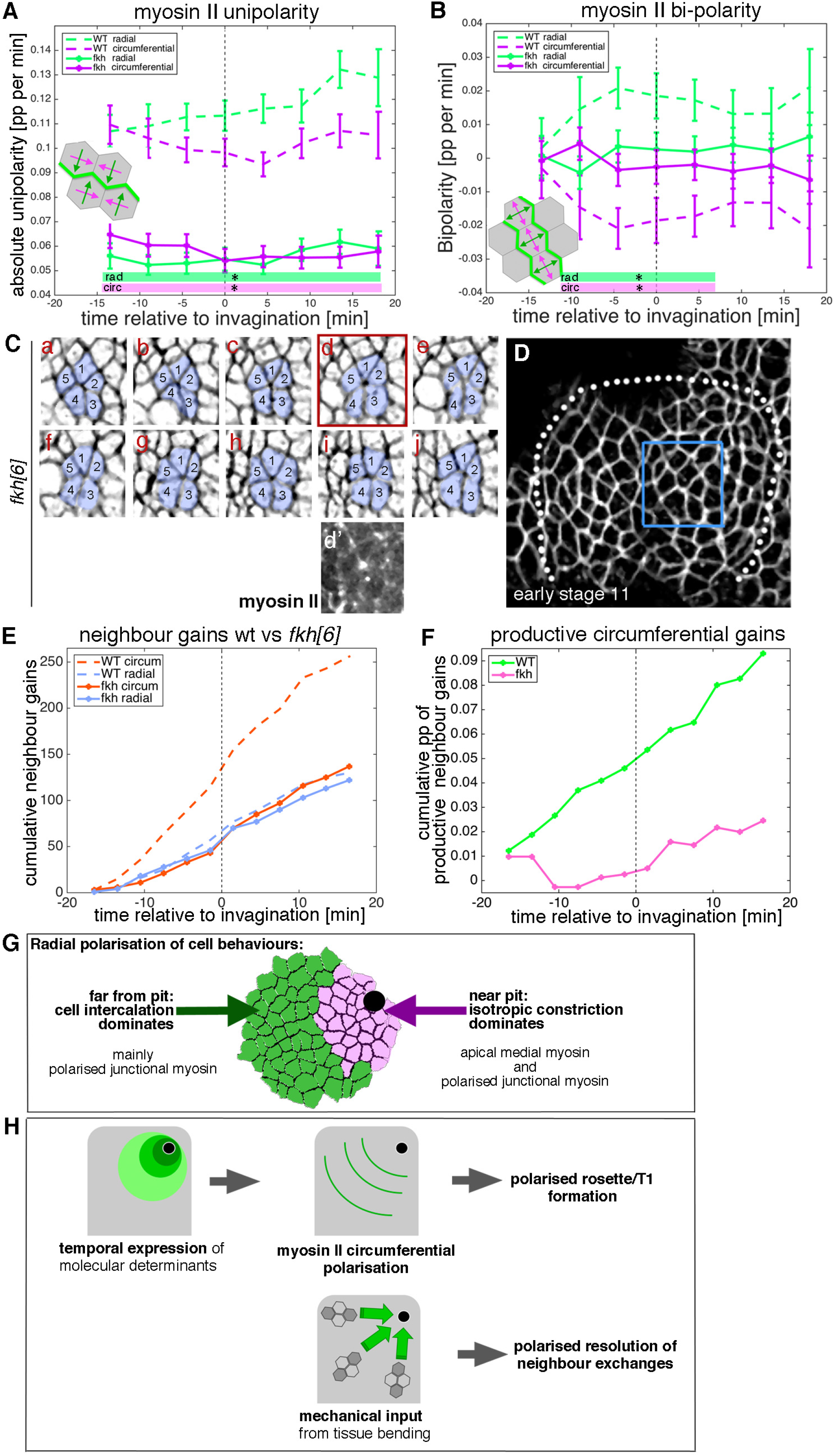
Analysis of intercalation behaviour in fkh mutants. **A,B** Analysis of myosin II unipolarity (**A**) and bi-polarity (**B**) in *fkh[6]* mutant placodes compared to wild-type (as shown in Figure 6, see also Suppl. Movie 6 and 7). Overall myosin II levels in *fkh[6]* mutant placodes are lower (not shown) and both unipolarity and bi-polarity are decreased, with bi-polarity near zero (**B**, solid curves). Error bars show intra-embryo variation of 5 embryo movies for *fkh[6]* and 4 embryo movies for wt. Statistical significance at p<0.05 using a mixed-effect model is indicated as shaded boxed at the bottom of the panels: wt vs *fkh[6]* for the radial vector/circumferential enrichment (green, ‘rad’) and wt vs *fkh* for the circumferential vector/radial enrichment (purple, ‘circ’). **C-F** Intercalation and neighbour exchanges still occur in *fkh[6]* mutant placodes. **C** Sill pictures from a tracked and segmented movie of a *fkh[6]* mutant placode (labelled with membrane-RFP), frames (labelled **a-j**) are 1:55 min apart, the full placode view in **D** is from time point **a, d’** shows the myosin accumulation at the centre of the rosette structure observed in **d**. **E** In comparison to wt where circumferential neighbour gains dominate over radial ones (dashed lines), in the *fkh[6]* mutant placodes both occur with equal frequency (solid lines). This leads to nearly no productive neighbour gains in the mutant, significantly fewer than in the wt (**F**; Kolmogorov-Smirnov two sample test D=0.5833, P=0.0191). **G** The placode shows a clear radial polarisation of cell behaviours, with isotropic constriction dominating near the pit (likely driven by the strong apical-medial actomyosin accumulation in these cells (Booth et al., 2014)), with cell intercalations dominating away from the pit, where junctional myosin is highly polarised. **H** The temporal expression of molecular determinants leads to myosin polarisation at circumferential junctions, thereby polarising and driving T1 and rosette formation. Mechanical force generated through the pulling of the invaginating pit polarises the resolution of intercalation events. Both the tissue-wide polarisation of myosin II and the polarisation of resolution are lost in the absence of Fkh.

Therefore, although at the tissue and cell shape level *fkh[6]* mutant placodes appear near static, the close analysis of individual events revealed a dynamic but unpolarised behaviour, with intercalation occurring, probably driven by the remnant myosin II polarisation, but with no polarised resolutions.

## Discussion

Morphogenesis sculpts many differently shaped tissues and structures during embryogenesis. A core set of molecular factors that are the actual morphogenetic effectors is repeatedly used, such as actomyosin allowing contractility or cell-cell adhesion components allowing coordination and mechanical propagation of cell behaviours across tissues. Such core morphogenetic effectors or effector modules are used reiteratively in different tissues and at different times, whereas the activity of upstream activating gene regulatory networks leading to tissue identity, but also initial tissue geometry and mechanical constraints, are highly tissue-specific.

During tube formation of the salivary glands in the fly embryo, we observe a clear tissue-level radial organisation of both cell behaviours and molecular factors driving these, with the off-centre located invaginating pit as the organising focal point (Fig. 8G,H). The cell behaviours of cell shape change and cell intercalation that we identified within the apical domain have previously been shown to drive other morphogenetic processes (Butler et al., 2009; Collinet et al., 2015; Lee and Harland, 2007; Lye et al., 2015; Martin and Goldstein, 2014; Martin et al., 2009; Plageman et al., 2011; Rauzi et al., 2010), but in our system, they are utilised within a radial coordinate system, thereby driving the formation of a narrow tube of epithelial cells from a round placode primordium. Interestingly, even the molecular factors that are involved in these behaviours in other contexts also contribute in the salivary gland placode, again being adjusted to create a radial/circumferential pattern. The main question to answer therefore is what allows the ‘twisting’ of a system that is earlier in embryogenesis used within an anterior-posterior/dorsal-ventrally polarised coordinate system into the radial coordinates observed here?

Our analysis suggests at least a two-fold contribution. A number of factors, both transcription factors as well as potential morphogenetic effectors, are expressed in the salivary gland placode in a temporally shifting pattern, initially restricted to the location of the forming invagination pit, and then slowly expanding in expression across the whole placode (Fig. 8H). Examples include the expression of the transcription factors Fkh (Suppl.Fig. 8A-C), Hkb and Fog, but also of factors such as the polarity factor and homophilic interactor Crumbs (Suppl. Fig. 8D’, H”, N), the nonreceptor tyrosine kinase Btk29A/Tec29 (Suppl.Fig.8 H’), the dynein-associated protein Klarsicht and the Toll-like receptor 18-wheeler (Chandrasekaran and Beckendorf, 2005; Kolesnikov and Beckendorf, 2007; Myat and Andrew, 2000b; Myat and Andrew, 2002; Panzer et al., 1992). Which of these or other factors expressed in a similar pattern leads to the radial tissue-polarisation of for instance myosin II and Bazooka across the placode is as yet unclear (Fig. 8H). Differing levels of Crumbs protein, with levels reducing with distance from the pit and a sharper step change in between secretory and duct primordium, could for instance assist myosin II accumulation in circumferentially-oriented junctions, as we have shown previously that step changes in Crumbs can pattern actomyosin accumulation (see schematic in Suppl.Fig.8N; (Röper, 2012; Thompson et al., 2013)).

In addition to this potential radial polarisation through upstream expression control, the positioning and invagination of the pit is another factor underlying the radial polarisation of behaviours, with for instance the intercalation strain rate increasing with the onset of invagination (Fig. 2E and Suppl.Fig.2C). In the *fkh* mutant no pit is specified and, over the time period analysed here, no invagination or tissue-bending is observed anywhere across the placode, and at the tissue-level, polarisation of myosin II is lost. Nonetheless, in individual rosette formation events (Fig. 8C) and in small regions of the placode (Suppl. Fig.8L-M) myosin II polarisation remains and neighbour exchanges occur at a rate comparable to the control but fail to contribute to an overall tissue change during early stages. In the *fkh* mutant several of the factors that in the wild-type show a temporal radial expression shift originating from the pit in the corner of the placode are now either homogeneously expressed across the placode (Btk29A, Suppl.Fig.8I,I’) or at higher levels in the centre of the placode (Crumbs, Suppl.Fig.8E’, I”), thereby most likely affecting the polarisation of donwstream factors. The circular topology of a flat epithelial primordium with tube budding from an off-centre location is not unique to the salivary gland placode but for instance also seen in the posterior spiracles in the *Drosophila* embryo (Simoes et al., 2006) as well as during dorsal appendage formation in the germline (Claret et al., 2014), so it will be interesting to determine in the future whether other aspects of the radial polarisation are in common.

Not only the fully invaginated epithelial tube of the salivary glands is a 3D structure, but already the epithelial primordium of the placode, though flat at this stage, should be considered as a 3D rather than a 2D entity, due to the apical-basal extension of the epithelial cells. Supporting this notion, our strain rate analysis in depth has revealed many interesting features. It is still quite unclear whether epithelial morphogenetic behaviours are initiated apically and propagated basally and in how much there is an active contribution through events initiated at lateral or basal sides. A few recent reports suggest that not all is apically initiated (Monier et al., 2015; Sun et al., 2017), but whether this is a general principle or highly tissue-specific is unclear. It is also unclear how any cell behaviour, whether initiated apically or basally, is communicated and propagated across the length of the cell. In many morphogenetic processes including the tube budding from the salivary gland placode, actomyosin and other morphogenetic effectors are concentrated within the apical junctional domain. Our data at depth reveal that there is close coordination of intercalation between apical and mid-basal levels, and that the dynamic changes in interleaving and wedging strongly support an active apical mechanism that is followed further basally.

Non-equilibrium 3D cell geometries in a flat epithelium, such as those caused by the wedging, interleaving and tilting analysed here, evolve in revealing patterns during early placode morphogenesis. The appearance and progressive increase in these out-of-equilibrium geometries in the placode precede the start of 3D pit invagination by more than 10 minutes. This could suggest that during this phase a pre-pattern of tension could build up across the placode that would make the pit invagination more efficient, once initiated. Some of the cell behaviours we observe and the resulting complex 3D shapes might then be the non-intuitive results of the balance of forces in the changing mechanical context of the placode. This will be possible to test experimentally by interference with certain behaviours or through ectopic induction of others, and can also be tested in silico in the future.

In summary, the radial patterning of the placode is key for both the radial-circumferential polarisation of the actomyosin cytoskeleton as well as for the specification of the first point of tissue-bending. As we have previously shown, in addition to the polarised junctional accumulation of myosin, an apical-medial pool of myosin accumulates most strongly in the cells near to the pit and drives the apical constriction of cells (Booth et al., 2014). This additional radial patterning of different actomyosin pools might well underlie the differing cell behaviours of cells near to and far from the pit (Fig.8G), with circumferentially enriched junctional myosin driving the initiation of intercalation events, be it T1s or rosettes, and medial myosin driving strong isotropic constriction and tissue bending of the invagination point. Once the tissue bending has initiated, the mechanical force of the invagination orients the polarised resolution of intercalation events (Fig.8H). Such mechanical influence on the orientation of intercalation is reminiscent of germband extension, where the pulling from the invagination of another neighbouring tissue, the posterior midgut, influences the intercalation (Collinet et al., 2015; Lye et al., 2015). In the salivary gland placode, radial patterning and mechanical input both originate from within the same tissue, and it will be interesting to explore how conserved such organisation is across other tube-forming tissue primordia.

## Supplemental Movies

***Suppl. Movie 1. Example movie of early salivary gland placode morphogenesis in 3D.***

Embryo of the genotype *Scribble-GFP/fkhGal4::UAS-palmYFP* as shown if Fig.1C and Fig.3A. Time stamp indicates time before and after initiation of tissue bending at t=0. Scale bar 20μm.

***Suppl. Movie 2. Example movies of Tissue strain rate tensor (SRT), Cell Shape SRT and Intercalation SRT.***

Local strain rates are extracted from segmented and tracked movies. Tissue shape change (left), cell shape change (centre) and intercalation (right) are shown. Red vectors represent contraction and blue vectors represent expansion. Vectors are projected onto a radial coordinate system. Time stamp indicates time before and after initiation of tissue bending at t=0. See also Figure 2.

***Suppl. Movie 3. Example of matching cells through depth.***

Apical cell identities matched to basal cell identities are shown within the placode. Stamp refers to the depth in the tissue. Scale bar 20μm. See also Figure 4.

***Suppl. Movie 4. Example movie of apical rosette formation/resolution.***

Embryo of the genotype *sqh[AX3]; sqh::sqhGFP42,UbiRFP-CAAX,* only the UbiRFP label is shown. A group of cells going through rosette formation/resolution is highlighted, still of the movie are shown in Fig.5E. Time stamp indicates time before and after initiation of tissue bending at t=0. Scale bar 20μm.

***Suppl. Movie 5. Example movie of cell shape and myosin II localisation in a control embryo.***

Embryo of the genotype *sqh[AX3]; sqh::sqhGFP42,UbiRFP-CAAX* used for the myosin II uni- and bi-polarity quantifications as shown in Fig.6. Time stamp indicates time before and after initiation of tissue bending at t=0. Scale bar 20μm.

***Suppl. Movie 6. Example movie of automatic junctional myosin II quantification.***

Myosin II uni-polarity vectors (left) and bi-polarity vectors (right) are shown. Junctions are colour coded according to their myosin II intensity levels as shown if Fig. 6C’ (middle). Time stamp indicates time before and after initiation of tissue bending at t=0. Scale bar 20μm. See also Figure 6.

***Suppl. Movie 7. Example movie of cell shape and myosin II localisation in a fkh[6] mutant embryo.***

Embryo of the genotype *sqh::sqhGFP42,UbiRFP-CAAX; fkh[6]* used for the myosin II uni- and bi-polarity quantifications as shown in Fig.8. Time stamp indicates time before and after initiation of tissue bending at t=0. Scale bar 20μm.

## Materials and Methods

### Fly stocks and husbandry

The following transgenic fly lines were used: *sqhAX3; sqh::sqhGFP42* (Royou et al., 2004) and *fkhGal4* (Henderson and Andrew, 2000; Zhou et al., 2001) [kind gift of Debbie Andrew]; *Scribble-GFP (DGRC Kyoto), UAS-palmYFP* (generated from membrane Brainbow; (Hampel et al., 2011)), *y*^*1*^ *w* cv*^*1*^ *sqh*^*AX3*^*; P{w^+mC^=sqh-GFP.RLC}C-42 M{w*^*+mC*^*=Ubi-TagRFP-T-CAAX}ZH-22A* (Kyoto DGRC Number 109822, referred to as *sqh*^*AX3*^*;sqhGFP; UbiRFP); P{w+mC=sqh-GFP.RLC}C-42 M{w+mC=Ubi-TagRFP-T-CAAX}ZH-22A; fkh[6]/TM3 Sb Twi-Gal4::UAS-GFP (fkh[6]* allele from Bloomington). See Table 1 for details of genotypes used for individual figure panels.

### Embryo Immunofluorescence Labelling, Confocal, and Time-lapse

Embryos were collected on apple juice-agar plates and processed for immunofluorescence using standard procedures. Briefly, embryos were dechorionated in 50% bleach, fixed in 4% formaldehyde, and stained with phalloidin or primary and secondary antibodies in PBT (PBS plus 0.5% bovine serum albumin and 0.3% Triton X-100). anti-Crumbs and anti-E-Cadherin antibodies were obtained from the Developmental Studies Hybridoma Bank at the University of Iowa; anti-Baz was a gift from Andreas Wodarz (Wodarz et al., 1999); anti-Btk/Tec29 was a kindly provided by Georgia Tsikala (Tsikala et al., 2014); anti-Fkh was a gift from Herbert Jäckle (Weigel et al., 1989). Secondary antibodies used were Alexa Fluor 488/Fluor 549/Fluor 649 coupled (Molecular Probes) and Cy3 and Cy5 coupled (Jackson ImmunoResearch Laboratories). Samples were embedded in Vectashield (Vectorlabs).

Images of fixed samples were acquired on an Olympus FluoView 1200 or a Zeiss 780 Confocal Laser scanning system as z-stacks to cover the whole apical surface of cells in the placode. Z-stack projections were assembled in ImageJ or Imaris (Bitplane), 3D rendering was performed in Imaris.

For live time-lapse experiments embryos from [*Scribble-GFP,UAS-palmYFP fkhGal4*], [*sqh^AX3^; sqhGFP,UbiRFP*] or [*sqhGFP,UbiRFP; fkh[6]*] were dechorionated in 50% bleach and extensively rinsed in water. Embryos were manually aligned and attached to heptane-glue coated coverslips and mounted on custom-made metal slides; embryos were covered using halocarbon oil 27 (Sigma) and viability after imaging after 24h was controlled prior to further data analysis. Time-lapse sequences were imaged under a 40x/1.3NA oil objective on an inverted Zeiss 780 Laser scanning system, acquiring z-stacks every 0.8-2.6 minutes with a typical voxel xyz size of 0.22 x 0.22 x 1 μm. Z-stack projections to generate movies in Supplementary Material were assembled in ImageJ or Imaris. The absence of fluorescent *Twi-Gal4::UAS-GFP* was used to identify homozygous *fkh[6]* mutant embryos. During the early stages of salivary gland placode morphogenesis analysed here, *fkh[6]* mutants showed no reduction in cell number or initiation of apoptosis (data not shown). The membrane channel images from time-lapse experiments were denoised using *nd-safir* software (Boulanger et al., 2010).

### Cell segmentation and tracking

Cell tracking was performed using custom software written in IDL (Blanchard et al., 2009; Booth et al., 2014). First, the curved surface of the embryonic epithelium was located by draping a ‘blanket’ down onto all image volumes over time, where the pixel-detailed blanket was caught by, and remained on top of binarised cortical fluorescence signal. Different quasi-2D image layers were then extracted from image volumes at specified depths from the surface blanket. We took image layers at 1-3 μm and 7-8 μm for the apical and mid-basal depths, respectively. Image layers were local projections of 1-3 z-depths, with median, top-hat or high/low frequency filters applied as necessary to optimse subsequent cell tracking.

Cells in image layers at these two depths were segmented using an adaptive watershedding algorithm as they were simultaneously linked in time. Manual correction of segmented cell outlines was performed for all fixed and time-lapse data. Tracked cells were subjected to various quality filters (lineage length, area, aspect ratio, relative velocity) so that incorrectly tracked cells were eliminated prior to further analysis. The number of embryos analysed can be found in Table 1 (and see Supplementary Fig. 2 (WT apical), Supplementary Fig. 3 (WT basal) and Supplementary Fig. 7 (fkh)).

### Mobile radial coordinate system for the salivary placode

WT movies were aligned in time using as t=0 mins the frame just before the first sign of invagination of cell apices at the future tube pit was evident. *fkh[6]* mutants were aligned using as a reference of embryo development the level of invagination of the tracheal pits that are not affected in the fkh[6] mutant as well as other morphological markers such as appearance and depth of segmental grooves in the embryo. Cells belonging to the salivary placode (without the future duct cells that comprise the two most ventral rows of cells in the primordium) were then manually outlined at t=0 mins using the surrounding myosin II cable as a guide and ramified forwards and backwards in time. Only cells of the salivary placode were included in subsequent analyses.

At t=0 mins, the centre of the future tube pit was specified manually as the origin of a radial coordinate system, with radial distance (in μm) increasing away from the pit (e.g. Fig. 1G). Circumferential angle was set to zero towards Posterior, proceeding anti-clockwise for the placode on the left-hand side of the embryo, and clockwise for the placode on the right so that data collected from different sides could be overlaid.

The radial coordinate system was ‘mobile’, in the sense that its origin tracked the centre of the pit, forwards and backwards in time, as the placode translated within the field of view due to embryo movement or to on-going morphogenesis.

### Morphogenetic strain rate analysis

Detailed spatial patterns of the rates of deformation across the placode and over time quantify the outcome of active stresses, viscoelastic material properties and frictions both from within and outside the placode. We quantified strain (deformation) rates over small spatio-temporal domains composed of a focal cell and one corona of immediate neighbours over a ~5 min interval ((Blanchard et al., 2009) and reviewed in (Blanchard, 2017)). On such 2D domains, strain rates are captured elliptically, as the strain rate in the orientation of greatest absolute strain rate, with a second strain rate perpendicular to this (Fig. 2B).

For the early morphogenesis of the salivary gland placode, in which there is no cell division or gain/loss of cells from the epithelium, three types of strain rate can be calculated. First, total tissue strain rates are calculated for all local domains using the relative movements of cell centroids, extracted from automated cell tracking. This captures the net effect of cell shape changes and cell rearrangements within the tissue, but these can also be separated out. Second, domain cell shape strain rates are calculated by approximating each cell with its best-fit ellipse and then finding the best mapping of a cell’s elliptical shape to its shape in the subsequent time point, and averaging over the cells of the domain. Third, intercalation strain rates that capture the continuous process of cells in a domain sliding past each other in a particular orientation, is calculated as the difference between the total tissue strain rates and the cell shape strain rates of cells. Strain rates were calculated using custom software written in IDL (code provided in (Blanchard et al., 2009) or by email from G.B.B.).

The three types of elliptical strain rate were projected onto our radial coordinate system (see Fig. 1F), so that we could analyse radial and circumferential contributions. Strain rates in units of proportional size change per minute can easily be averaged across space or accumulated over time. We present instantaneous strain rates over time for spatial subsets of cells in the placode, and cumulative strain rates for the same regions over time. These plots were made from exported data using MATLAB R2014b.

We calculated strain rates in layers at two depths for WT placodes. Because we applied a radial coordinate system originating in the centre of the future tube pit to both depths, and used the same reference t=0 mins, we were able to compare strain rates at the same spatio-temporal locations on the placode between the depths.

### Matching cells between apical and mid-basal layers

We also wanted to characterize the 3D geometries of cells within small domains, using the cell shapes and arrangements in the two depths we had tracked. To do this, we needed to correctly match cells between apical and mid-basal depths. We manually seeded 3-5 apico-basal cell matches per placode, then used an automated method to fill out the remaining cell matches across the placode and over time.

We did this by sequentially looking at all unmatched apical cells that were next to one or more matched cell. The location of the unmatched basal centroid was predicted by adding the vector between matched and unmatched apical cell centroids to the matched basal centroid. The nearest actual basal centroid to this predicted basal centroid location was chosen as the match, on condition that the prediction accuracy was within 0.25 of the apical centroid distance. Information from multiple matched neighbours was used, if available, improving the basal centroid prediction. Progressively matching cells out from known matched cells filled out placodes for all embryos. We visually checked apical-basal matches for each embryo in movies of an overlay of apical and basal cell shapes with apico-basal centroid connections drawn (Suppl. Movie 3).

### z-strain rates: 3D domain geometries

Having matched cells between apical and mid-basal layers, we have access to information about approximate 3D cell shapes. For single cells we can measure how wedged the cell shape is in any orientation and how tilted is the apical centroid to basal centroid ‘in-line’ relative to the surface normal (Suppl. Fig. 4A). In particular, we are interested in the amount of wedging and tilt in our radial and circumferential placode orientations (Fig. 4A). However, an important aspect is missing which is that cells can be arranged differently apically versus basally, that is they can be to a greater or less extent interleaved. The degree of interleaving is a multi-cell phenomenon, so we return to our small domains of a cell and its immediate neighbours. Within these small domains we want to quantify the amount of cell wedging, interleaving and cell tilt.

To correctly separate these quantities, we borrow from methods developed to separate out the additive contributions of wedging, interleaving and tilt that account for epithelial curvature (Deacon, 2012). Deacon shows that for small domains of epithelial cells, curvature across the domain is the sum of their wedging and interleaving, while cell tilt has no direct implication for curvature.

During our study period, salivary placodes have minimal curvature. We therefore simplify the problem to flat (uncurved) domains, uncurving any local curvature so that apical and basal cell outlines are flat and parallel. Usefully, we can then treat small domains in exactly the same way as we have done above to calculate strain rates, but instead of quantifying the rate of deformation over time, here we calculate rate of deformation in depth, or ‘z-strain rates’.

The cell shape strain rate is equivalent to the wedging z-strain rate (Fig. 4B) and the intercalation strain rate is the interleaving z-strain rate (Fig. 4D). Both of these add up to a total strain rate and a total z-strain rate (Suppl. Fig. 4C), respectively. Note that the total z-strain rate of a domain can be the result exclusively of wedging or of interleaving (as in Fig. 4B,D; as with temporal strain rates, Fig. 2A), but some combination of the two is more likely. Domain translation and rotation for temporal domains become domain tilt (Fig. 4F) and torsion in the 3D domain geometries, respectively. Domain torsion is very weak in spatial or temporal averages of the placode (data not shown).

The units of wedging and interleaving are proportional size change per μm in z, and tilt is simply a rate (xy μm / z μm). We use the convention that change is from basal to apical, so a bottle-shape would be negative cell wedging. The number of embryos analysed for 3D cell geometries can be found in Table 2.

### Fluorescence intensity quantifications of Myosin and Bazooka

Embryos of the genotype *sqhAX3; sqh::sqhGFP42* (Royou et al., 2004) were labelled with either with anti-Bazooka and anti-DE-Cadherin or anti-DE-Cadherin and phalloidin to highlight cell membranes. For Fig. 5B, myosin II signal intensity of junctions oriented either circumferentially or radially from 5 placodes was quantified using Image J. For Fig. 5I fluoresce intensity of myosin II and Bazooka of junctions such as the ones indicated by arrows in Fig. 5H’, that were oriented either circumferentially or radially from 10 placodes was quantified using Image J. 3-pi wide lines were manually drawn along each junction. Intensity values were normalized to average fluorescence outside the placodes. We used a paired t-test for comparison of intensities within in the same junction, unpaired t-tests for comparison of circumferential myosin II vs radial myosin II and circumferential Baz vs radial Baz.

### Automated Myosin II quantification and polarity

Whereas previously we quantified apicomedial Myosin II (Booth et al., 2014), here we focused on junctional Myosin II. We extracted a quasi-2D layer image from the Myosin II channel at a depth that maximised the capture of the junctional Myosin. We background-subtracted the Myosin images and quantified the average intensity of Myosin along each cell-cell interface within the placode. To calculate the average intensity, we set the width of cell-cell interfaces as the cell edge pixels plus 2 pixels in a perpendicular direction either side. This captured the variable width of Myosin signal at interfaces.

We further summarised the uni- and bipolarity of Myosin for each cell. Methods to calculate Myosin II unipolarity and bipolarity are described in detail in (Tetley et al., 2016). Briefly, the average interface fluorescence intensity around each cell perimeter as a function of angle is treated as a periodic signal and decomposed using Fourier analysis. The amplitude component of period 2 corresponds to the strength of Myosin II bipolarity (equivalent to planar cell polarity) (Fig. 6D’). Similarly, the amplitude of period 1 is attributed to myosin unipolarity (junctional enrichment in a particular interface) (Fig. 6D”). The extracted phase of period 1 and 2 (bi/unipolarity) represent the orientation of cell polarity. We projected both polarities onto our radial coordinate system. Both polarity amplitudes are expressed as a proportion of the mean cell perimeter fluorescence. Cells at the border of the placode neighbouring the supra-cellular actomyosin cable were excluded from the analysis. The number of embryos analysed can be found in Table 2.

### Neighbour exchange analysis

We used changes in neighbour connectivity in our tracked cell data to identify neighbour exchange events (T1 processes). Neighbour exchange events were defined by the identity of the pair of cells that lost connectivity in *t* and the pair that gained connectivity at *t+1*. The orientation of gain we defined as the orientation of the centroid-centroid line of the gaining pair at *t+1.* We further classified gains as either radially or circumferentially oriented, depending on which the gain axis was most closely aligned to locally. We did not distinguish between solitary T1 s and T1 s involved in rosette-like structures.

From visual inspection, we knew that some T1s were subsequently reversed, so we characterised not only the total number of gains in each orientation but also the net gain in the circumferential axis, by subtracting the number of radial gains. Furthermore, when comparing embryos and genotypes, we controlled for differences in numbers of tracked cells by expressing the net circumferential gain per time step as a proportion of half of the total number of tracked cell-cell interfaces in that time step. We accumulated numbers of gains, net gains, and proportional rate of gain over time for WT (Fig. 5C) and *fkh* (Fig. 8E,F) embryos. Two sample Kolmogorov-Smirnov test was used to determine significance at p<0.05 for data in Fig. 8F.

### Statistical analysis of time-lapse data

Statistical tests to determine significance of data shown are indicated in the figure legends. Significance in time-lapse movies was calculated for bins of 4.5 minutes using a mixed-effects model implemented in R (‘lmer4’ package as in (Butler et al., 2009; Lye et al., 2015)) using variation between embryos as random effects and a significance threshold of p<0.05.

Error bars in time-lapse plots show an indicative confidence interval of the mean, calculated as the mean of within-embryo variances. The between-embryo variation is not depicted, even though both are accounted for in the mixed effects tests.

## Acknowledgements

The authors would like to thank the following people; for reagents and fly stocks: Debbie Andrew, Andreas Wodarz, Georgia Tsikala, Herbert Jäckle; for help with image analysis: Jerôme Boulanger.

Work in the Röper lab is supported by the Medical Research Council (file reference number U105178780). GBB was supported by grant no. 15.23(k) from the Isaac Newton Trust, by Wellcome Trust grant no. 100329/Z/12/Z awarded to William Harris and Biotechnology and Biological Sciences Research Council Standard Grant BB/J010278/1 to Richard Adams and Bénédicte Sanson.

